# Protein Dynamics Underlies Strong Temperature Dependence of Heat Receptors

**DOI:** 10.1101/2024.11.04.621882

**Authors:** Andrew Njagi Mugo, Ryan Chou, Feng Qin

## Abstract

Ion channels are generally allosteric proteins, involving specialized stimulus sensor domains conformationally linked to the gate to drive channel opening. Temperature receptors are a group of ion channels from the transient receptor potential (TRP) family. They exhibit an unprecedentedly strong temperature dependence and are responsible for temperature sensing in mammals. Despite intensive studies, however, the nature of the temperature sensor domain in these channels remains elusive. By direct calorimetry of TRPV1 proteins, we have recently provided a proof of principle that temperature sensing by ion channels may diverge from the conventional allosterity theory; rather it is intimately linked to inherent thermal instability of channel proteins. Here we tackle the generality of the hypothesis and provide key molecular evidences on the coupling of thermal transitions in the channels. We show that while wild-type channels possess a single concerted thermal transition peak, the chimera, in which strong temperature dependence becomes disrupted, results in multi-transition peaks, and the activation enthalpies are accordingly reduced. The data show that the coupling with protein unfolding drives up the energy barrier of activation, leading to a strong temperature dependence of opening. Furthermore, we pinpoint the proximal N-terminus of the channels as a linchpin in coalescing different parts of the channels into concerted activation. Thus, we suggest that coupled interaction networks in proteins underlie the strong temperature dependence of temperature receptors.

**Significance:** Decoding receptor mechanisms requires understanding receptor activation at molecular levels. Whereas structural studies can unravel critical residues participating in activation, functional measurements are ultimately needed to pinpoint their mechanistic roles. Temperature receptors are gateways to thermosensation and pain. Despite intensive studies, how they detect temperature remains elusive. Here, by directly measuring heat flow in the most temperature-sensitive, high-threshold noxious heat receptor TRPV2, we show that channel activation is accompanied with a heat uptake sufficient to induce protein unfolding. We present molecular evidence that heat activation and unfolding are coupled, and propose a new mechanism based on concerted activation of different parts of channels to drive up temperature sensitivity. Our findings provide a mechanistic framework for understanding thermal biological processes.

## Introduction

Ion channels are versatile molecular sensors for detecting a wide array of cellular and environmental cues. To be able to detect a small change in stimuli, many of them evolve an exquisite sensitivity. The mechanisms underlying this sensitivity for many channels are broadly understood at least in principle. Voltage-gated channels, for instance, utilize localized lipid charges to maximize their responsiveness to changes in membrane potential, while ligand-gated channels exploit multimeric binding to enhance agonist sensitivity ^1^. Temperature receptors, however, stand a notable exception to this understanding. Despite intense studies, the origins of their pronounced temperature sensitivity remain largely elusive in these channels.

The most extensively studied temperature receptors belong to the transient receptor potential (TRP) superfamily ^2,3^. Out of the 28 TRP channels, eleven are directly activated by specific temperatures and have been implicated in temperature sensing by mammals. The majority of them are heat-sensitive and fall into the vanilloid and melastatin subfamilies (TRPV^4-8^, TRPM ^9-14^), though the cold receptor TRPM8 is also a member of the TRPM family ^15,16^. One characteristic feature of thermoreceptive neurons is their ability to produce graded-responses to temperature changes ^17^, which dictate that multiple temperature receptors exist. Consistently, thermal TRPs are tuned to detect specific temperature ranges, such as hot, cold, or warm. The Q_10_ commonly denote the rate of activity increase of TRP channels over a 10°C temperature rise. Thermal TRPs often exhibit Q_10_ values significantly higher than those of other channels (e.g. > 20 ^18^ for TRPV1). Some, such as the high-threshold noxious heat receptor TRPV2, even reaches Q_10_ ∼ 100 ^19^.

The exquisite temperature sensitivity of thermal TRPs has inspired extensive structure-functional studies, aimed to locate novel temperature sensors within the channels. While studies affirm the intrinsic temperature sensitivity of these channels ^20-23^, they stop short of converging to a consensus location. Instead, the identified molecular sites are seemingly distributed throughout the entire channels ^24-42^. These include the pore region, the voltage-sensing domain (S1-S4), the membrane proximal N-terminus or the pre-S1 linker, the ankyrin repeats, the C-terminus, etc. More recent Cryo-EM studies further added to the list of additional moving parts of the channels and lipids ^35-37^. One appealing theory links the temperature dependence of the channels to changes in heat capacity, a phenomenon commonly associated with the solvation of hydrocarbons ^43-46^. Nonetheless, a lack of clear endothermic or exothermic features at many of the identified molecular sites has left the precise mechanisms by which thermal TRPs sense temperature still an open question.

Irrespective of the molecular basis, thermodynamics dictates that the temperature-induced channel opening is associated with a large enthalpy change (ΔH). The opening of TRPV1, for example, was estimated to involve an enthalpy change of approximately 100 kcal/mol ^47^. Conventional patch-clamp estimates do not take into account the energy dissipated through allosteric coupling, suggesting that the actual energy absorbed by the channels at the onset would be substantially larger. Our recent work using differential scanning calorimetry (DSC) provided a direct measure of TRPV1’s energy consumption ^48^, showing that the heat uptake during channel activation far exceeds patch-clamp estimates. This substantial energy uptake corresponds to the range typically associated with protein unfolding, indicating that the heat-evoked opening is accompanied with partial protein unfolding. Thus, the DSC supports a model that the coupling of channel activation with protein unfolding underlies the strong temperature dependence of the channel. But the model has only been elucidated for TRPV1.

Here we explore the generality of the unfolding mechanism, focusing particularly on the molecular basis of the coupling of thermal transitions in ion channels. We investigated the thermal transitions of TRPV2 and its chimera by DSC. We show that the diminished temperature dependence in the chimera correlates with reduced energy barriers of thermal transitions and disruption of cooperative interactions between component transitions. The results support the idea that concerted cross-domain activation in channels underlies their substantial energetic characteristics. Specifically, we identify the proximal N-terminus to be a crucial region for forging cross-domain interactions to coalesce different parts of the channels into concerted transitions. Based on the significance of the coupling interactions, we suggest that protein dynamics plays an essential role in temperature sensing by ion channels and possibly other biological thermal processes.

## Results

### 1. Thermal transitions in TRPV2

To detect mutagenic effects using DSC, we selected TRPV2, known for its high temperature-sensitivity, as a wild-type template ^19^. We reasoned that its large temperature dependence would provide an adequate leeway for mutations to substantially alter the DSC transition while still retaining a detectable residual thermal sensitivity. TRPV2 also has a higher temperature threshold of activation than TRPV1, further making it a good candidate for examining the subtype specificity of DSC measurements.

TRPV2 proteins were purified by affinity chromatography using the maltose binding protein (MBP) tag fused to the N-terminus of the channel. Electrophoresis of purified channel proteins ensured a single-band purity on SDS-PAGE gels (Fig. 1a). Size-exclusion chromatography verified the sample predominantly in a monodispersed form at the anticipated tetrameric size (Fig. 1b). For functional and DSC studies, we reconstituted the channel proteins into vesicles formed from soybean lipids. We generally kept the MBP tag during reconstitution for downstream quantification of protein concentrations in vesicles, using the melting energy of MBP as a caliper (see Methods). The MBP tag in these channels does not interfere with channel functions ^21^. Patch-clamp of the reconstituted channels showed their native sensitivities to both agonist and heat (Fig. 1c-e). In particular, the heat activation occurred in the range of 48°C – 53°C, close to that in native membranes, and the activity tended to be transient, lasting over a few heating pulses (each 100 ms) before fading away (Fig. 1e), a phenomenon referred to as rundown hereafter and also observed in TRPV1 ^48,49^.

**Figure 1.**
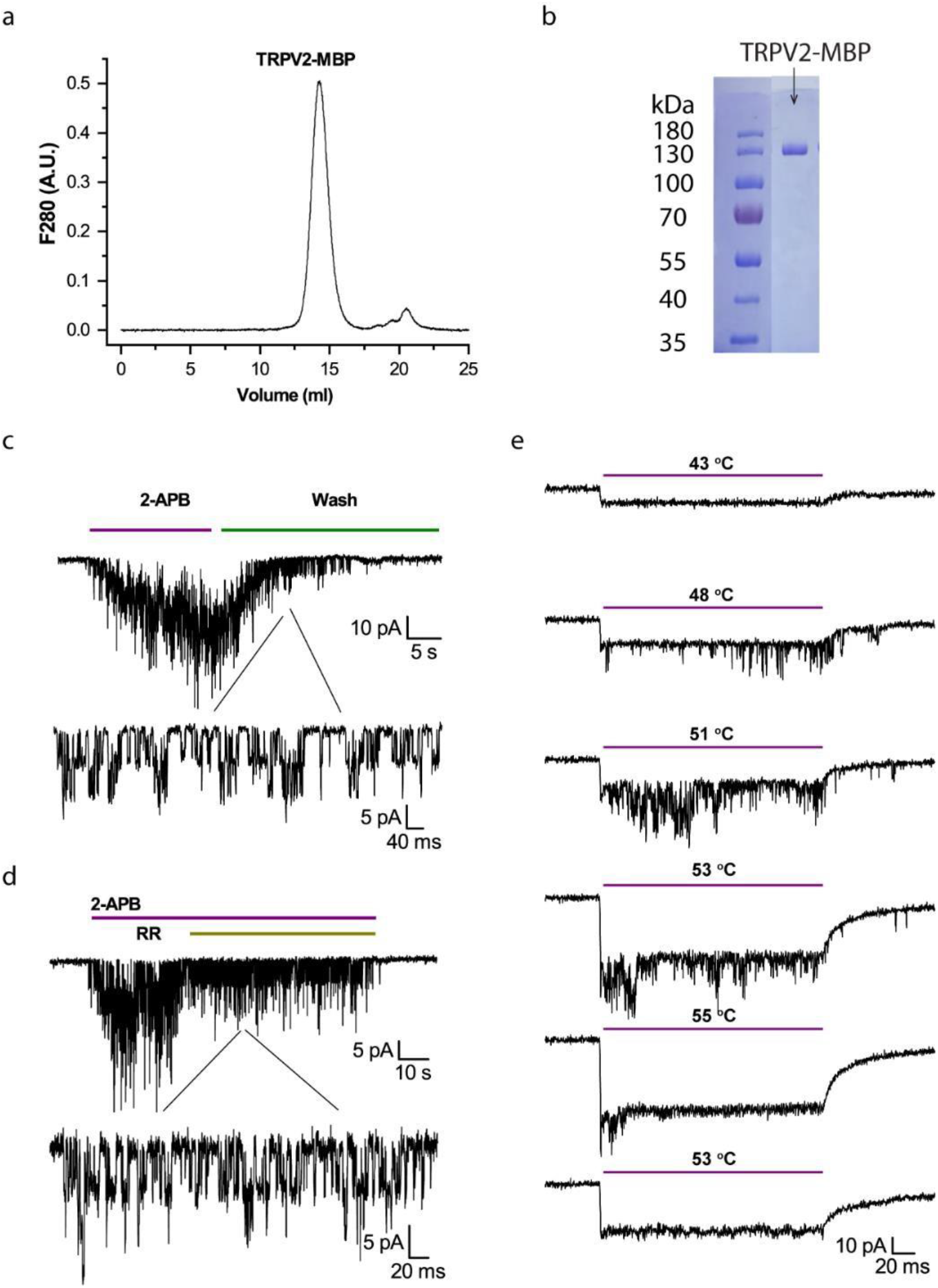
Characterization of purified and reconstituted TRPV2 proteins. **A)** Size-exclusion chromatograph of purified TRPV2-MBP proteins in a Superose 6 Increase 10/300 column. **B)** Electrophoresis profile of purified TRPV2-MBP proteins on Coomassie-blue-stained SDS-PAGE. **C)** Agonist responses of reconstituted TRPV2 in liposomes evoked by 2-APB (1mM). **D)** Inhibition of 2-APB responses by ruthenium red (RR, 10mM). **E)** Heat responses evoked by a series of temperature pulses (each 200 ms long). From top to bottom: Channel openings first occurred between 48°C – 53°C and then ran down over continued heating to 55°C. Reducing temperature in subsequence pulses did not recover the activity. The immediate current drop at the beginning of each temperature pulse was due to leakage of patch. Holding potential -60mV.

The DSC scan of the TRPV2-MBP proteins detected two thermal transition peaks riding on a slowly varying instrument baseline (Fig. 2a). The soybean lipids we used for reconstitution are free of phase transitions over a broad temperature range (5 °C -70 °C, ^48^). Thus, both transition peaks were attributable to the added channel proteins (TRPV2-MBP). Fig. 2b illustrates the isolated transition peaks (a.k.a. thermographs) after subtracting the baseline. Of the two transitions, the second (Fig. 2b, inset) coincided with the melting of the MBP protein ^50^, which was similarly observed in the DSC of TRPV1-MBP ^48^. Thus, we attributed it to the fusion tag MBP. Consistently, the first peak (Fig. 2b) occurred at temperatures corresponding to heat-evoked opening of TRPV2, and differed from the thermal transition of TRPV1 (Fig. 2c). The latter supports the subtype-specificity and functional correlation of the DSC measurement, leading us to attribute the first peak to the TRPV2 protein.

**Figure 2.**
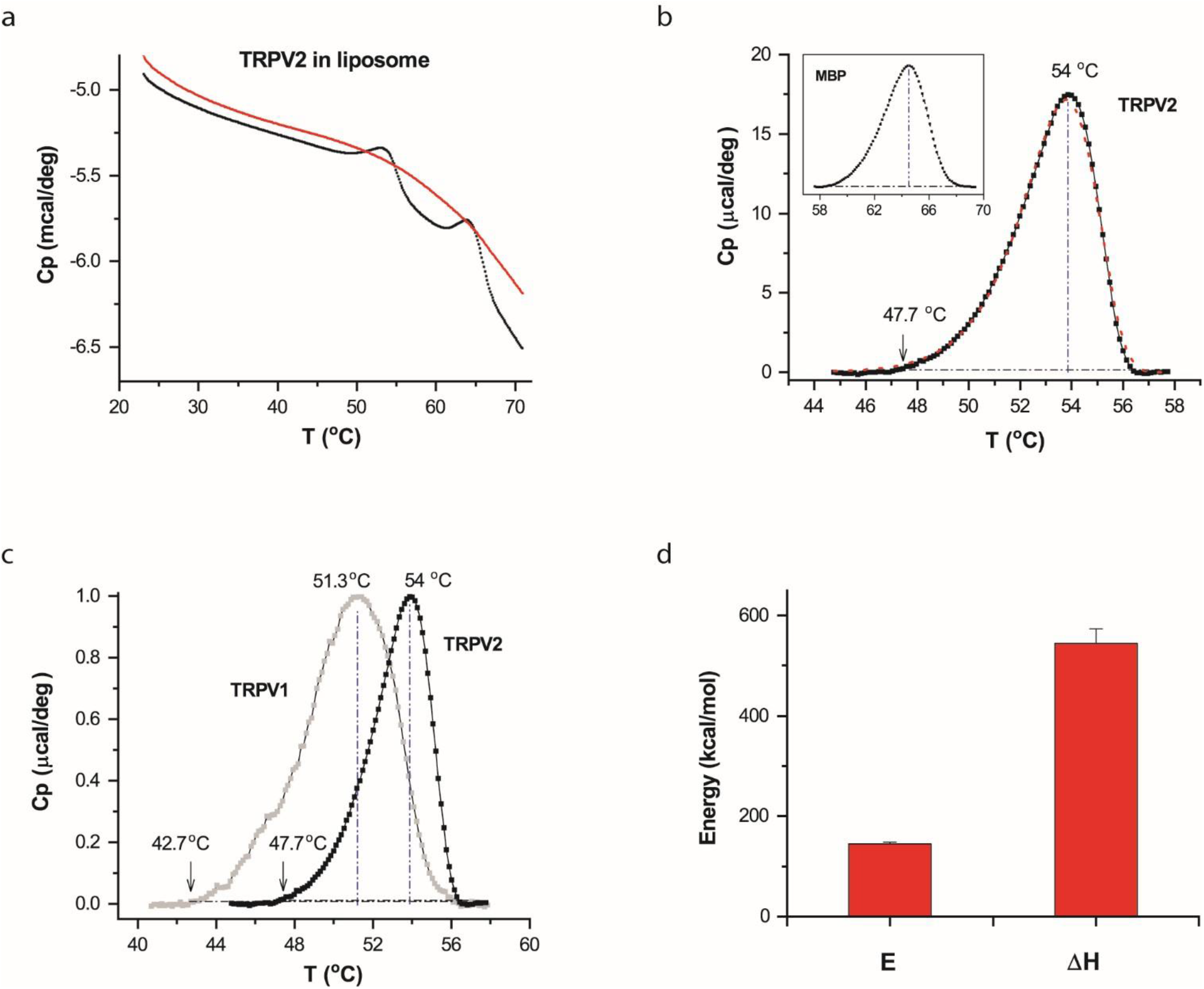
DSC transitions of TRPV2 in vesicles. **A)** Representative DSC scans of TRPV2 proteins reconstituted in liposomes. The trace in red corresponds to repeated scan, showing the irreversibility of the DSC transition of TRPV2. **B)** Thermograph of excess heat capacity of TRPV2 proteins (dotted black curve). The red curve represents the fit by a two-state model. The vertical line and the arrow indicated temperatures where the transition reached the peak or 1% of the peak amplitude. The inset shows the profile of the excess heat capacity of MBP fused to the channel protein. **C)** Comparison of DSC transitions of TRPV2 (solid black) and TRPV1 (grey), showing that the transitions of TRPV2 occurred at higher temperatures as expected from its higher heat activation temperature. The temperature thresholds of the DSC transitions were also in the range of the temperature threshold of the heat activation of TRPV2. **D)** Average plot of DSC enthalpy change (ΔH) and activation energy (E) of TRPV2 in liposomes (n=9). DSC scanning rate was 1.0 °C/min.

To quantify the thermographs, we fitted them to appropriate models using predicted excess heat capacity (S10). Fig. 2b illustrates the fit (red line) by a two-state model, A→B, parametrized by an end-state enthalpy change (ΔH) and an energy barrier height or activation enthalpy (E). The fitting resulted in ΔH=545±29 kcal/mol and E = 145±3.4 kcal/mol (n=9). For comparison with patch-clamp responses, we also characterized the temperature threshold of DSC transitions, which, measured at ∼5% of the peak, was on an average around 48 °C. Both the temperature threshold and the activation energy agreed well with patch-clamp measurements ^24,26,51^, consistent with the expectation that the channels are functionally active in DSC and that the opening by heat led the DSC transition. The ΔH between the closed and open states of TRPV2 remains unresolved by patch clamp; thus it is not possible to compare the measured ΔH with patch-clamp estimates. But the value (∼545 kcal/mol) was in the range of, though larger than, that of TRPV1 (∼400 kcal/mol ^48^), indicating that TRPV2 has an even larger heat-uptake capacity.

The ΔH of TRPV2 is considerably larger than the activation energy E, implying that substantial structure changes occurred after the transition state. Consistently, the value of ΔH, on a scale of 6.3 cal/g, was in the range of protein unfolding ^52^. Thus, the thermal transitions of TRPV2 also encompassed at least channel opening and (partial) protein unfolding, as observed in TRPV1. This multi-state complexity of the DSC transition is in accordance with the observation of functional rundown after initial opening in patch-clamp (Figs. 1e & S7). The DSC heating protocol proceeded at a slow rate; therefore we further examined whether the unfolding transition in DSC was related to prolonged heating. Specifically, we exploited more rapid heating of the sample in a thermocycler before loading it into DSC. As shown in Fig. 3a, when the sample was preheated to 54 °C (2 min) in a thermocycler, the DSC transition was largely diminished, indicating that the unfolding transition also occurred by the thermocycler preheating. Temperature controls in thermocyclers tend to be less precise as the heating duration becomes shorter. Nevertheless, shortening the apparent heating duration in the thermocycler to 30 s remained effective (Fig. 3b). Therefore, the experiments suggest that the unfolding transition of TRPV2 was intrinsic to the energy landscape of TRPV2 and independent of DSC protocols.

**Figure 3.**
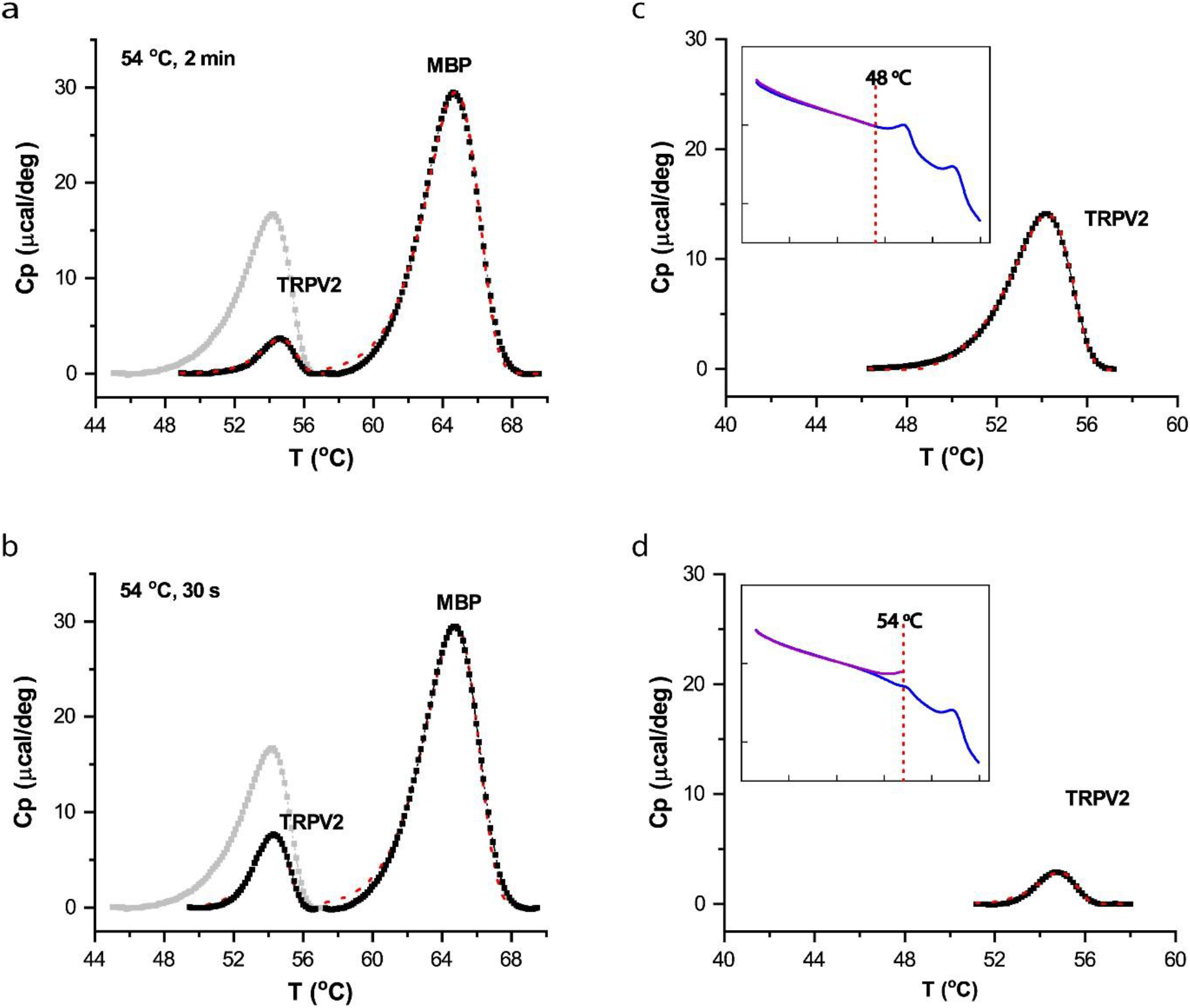
**A**,**B)** Independence of protein unfolding transition on DSC heating rate. Instead of being slowly heated up in DSC, samples were more rapidly preheated to target temperature (54 °C) in a thermocycler before loaded into DSC. Shortening heating duration did not prevent the unfolding transition. Panel A shows that a thermocycler heating for 2 min sufficed to significantly reduce DSC transition of TRPV2, while Panel B shows that even a 30-s treatment remained effective, causing significant unfolding of TRPV2 proteins. The thermograph of TRPV2 was displayed on a scale relative to the transition of the MBP tag. The grey trace shows control transition of TRPV2 (no prior heat treatment) for comparison. **C**,**D)** Stringent temperature dependence of DSC transition of TRPV2. Restricting initial DSC scan to a sub-transition temperature (48 °C) largely ameliorated irreversible loss of DSC transitions in repeated scans, indicating the temperature specificity of the transition and the independence on prolonged DSC heating (C). Accordingly, increasing scan temperature to 54 °C diminished the DSC transition in the repeated scan (D). Plotted are residual DSC transitions detected in the repeated scan, along with actual DSC scans (insets).

### 2. Concertedness of thermal transitions

Despite apparently involving multiple components (e.g. opening and unfolding), it is interesting that the DSC transition appeared as a single concerted peak to traverse a single activation energy barrier. One explanation is that the component transitions were mechanistically coupled so that the opening transition lowered the energy barrier of unfolding, so DSC detects only the initial energy barrier of opening. To test the hypothesis, we examined the propensity of component transitions to perturbations, particularly environmental changes. We reasoned that if the component transitions occurred independently and involved different parts of channel proteins, they could be differentially impacted by environmental changes. Fig. S1 shows the DSC transition of TRPV2 after the replacement of lipids by detergents. The channel in micelles remained strongly endothermic, with a comparable ΔH to that in lipids (Fig. S1c). However, the transition temperature was significantly shifted, from T_m_=54 °C in lipids to T_m_=50.5 °C in micelles (Fig. S1b). Importantly, DSC still detected a single concerted transition peak, indicating that the individual component transitions were shifted concomitantly, thus supporting that they were interconnected.

In a further experiment, we investigated whether limiting DSC scan temperature could prevent the unfolding transition, thereby separating it from the opening transition (Fig. 3c,d). When the final temperature in the initial scan was reduced to 48 °C, the subsequent full-temperature scan still detected the strong transition peak of TRPV2 proteins at 54 °C (Fig. 3c), suggesting that the channel proteins were stable irrespective of the slow heating up to 48 °C. On the other hand, increasing the temperature of the initial scan to ∼54 °C largely suppressed the DSC transition in the following scan (Fig. 3d). This indicates that the DSC transition in the channel had a strict temperature dependence, and was inducible by a submaximal opening temperature. The findings corroborate the inter-dependence of the opening and unfolding transitions ^48^.

### 3. Role of the proximal N-terminus

The coupling of channel opening to protein unfolding alluded to the existence of cooperative domain interactions linking the gate to other parts of the channel. We reasoned that such cross-domain interactions could forge a resistive force to the gate movement, thereby increasing the energy barrier of channel opening. Conversely, disrupting the interactions would allow the gate to move more freely and consequently reduce the temperature dependence. Guided by this rationale, we examined molecular perturbations in TRPV2 that can significantly alter its heat sensitivity, to draw molecular evidence on the coupling interactions.

The membrane proximal N-terminal domain (MPD) is a pre-S1 linker located between the intracellular ankyrin repeats and the transmembrane domains (TMD) (Fig. 4a). In the Cryo-EM structures of these channels, it interfaces with core parts of the channels such as TRP helix, ARD and TMD ^53,54^. And its perturbation has profound effects on the heat sensitivity of the channels ^26^. Thus, we examined whether the region pertains to the concerted DSC transition of TRPV2. We replaced the region in TRPV2 by the counterpart of TRPV1. The resultant chimera, TRPV2/V1 was similarly purified as wild-type TRPV2 (Fig. S2 a-b), and the reconstituted chimeric channels also remained functional (Fig. S2 c-e). Figs. 5b shows DSC thermographs of the chimera in vesicles. They were strikingly different from the wild-type transitions of TRPV2 or TRPV1. While the wild-type channels displayed a single transition peak, the chimera exhibited two segregated transition peaks. The larger peak occurred at higher temperatures and resembled more closely the wild-type TRPV2 transition (Fig. 4c, grey), but with noticeable changes in both peak steepness and transition temperature. The smaller peak preceded the larger one with a rather shallow slope (low activation energy E). These differences unravel a critical role of the region on the energy landscape of heat sensing by the wild-type channel.

**Figure 4.**
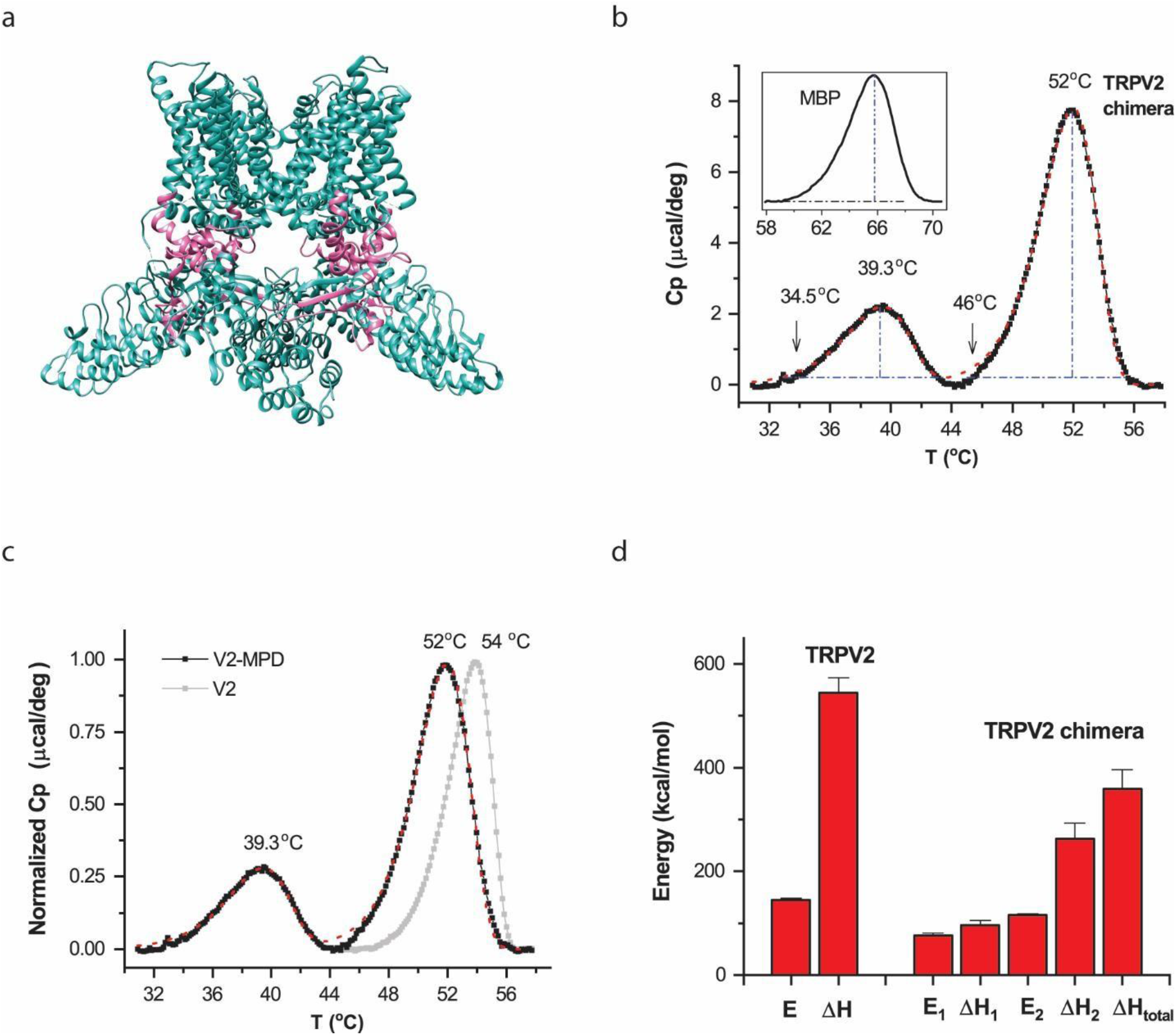
Thermal transitions of chimeric channel TRPV2/V1(357-434) in liposomes. **A)** Cartoon of the chimeric protein illustrating the location of the exchanged region (red color) in channels. The chimera contained swapping of the membrane proximal N-terminal domain (MPD) in TRPV2 by that of TRPV1. **B)** Thermograph of excess heat capacity. The chimera exhibited two split thermal transitions. The red line shows the fit by a three-state model (A → B → C). The arrow shows the temperature threshold at 1% of the peak amplitude. The inset shows the thermograph of fused MBP proteins. The melting temperature of MBP was ∼64 °C, which remained the same as when fused to the TRPV2 wild-type. Protein concentration was 0.53 mg/ml. **C)** Comparison of DSC transitions between the chimera and the wild-type channel. Swapping MPD fundamentally altered the wild-type transition profiles. **D)** Summary plot of energetics of chimera (n=8) versus wild-type TRPV2. E_1_ and ΔH_1_ are respectively the activation energy and the end-state enthalpy change for the first transition (A→B) while E_2_ and ΔH_2_ correspond to those of the second transition (B→C). ΔH_total_ represents the overall enthalpy change across all states. DSC scanning rate was 1.0 °C/min.

**Figure 5.**
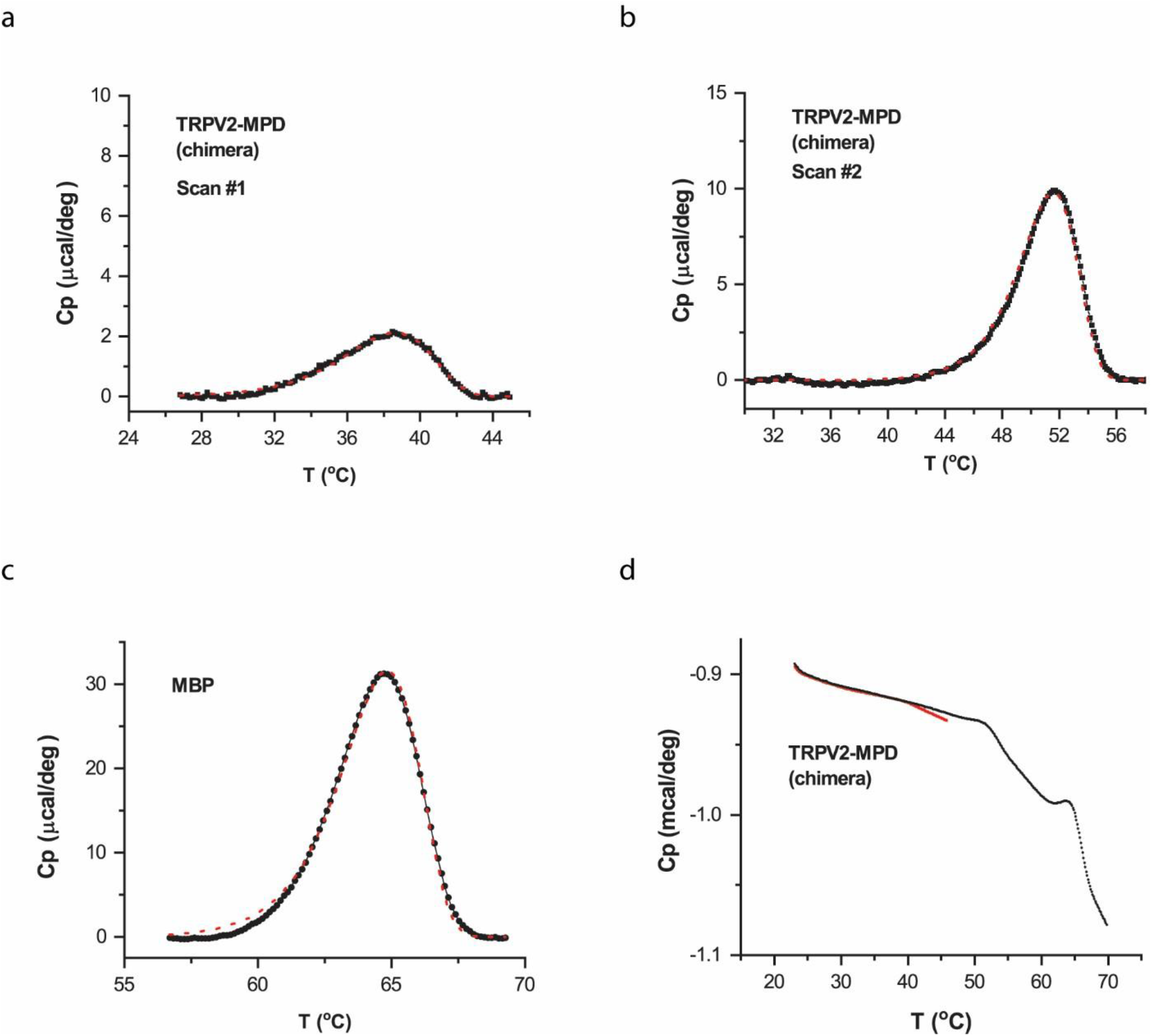
Irreversibility of thermal transitions in chimera. Reconstituted TRPV2/V1(357-434) chimeric proteins in vesicles were first scanned with final temperature up to 45 °C to manifest the first DSC transition peak while limiting the second high-temperature transition. The scan was then repeated to final temperature 70 °C. **A)** The minor, low-temperature DSC transition detected during the initial scan. **B, C)** The transitions occurring during the second scan. The predominant high-temperature transition peak remained, but the low-temperature peak that occurred between 32 °C and 44 °C during the first scan became unmeasurable (B). **D)** Representative DSC scans. Red for the initial scan and black for the repeat. The rate of temperature increase was 1 °C/min. The red dotted lines in (a)-(c) correspond to the best fits by an irreversible two- or three-state model.

To quantify the energetic changes, we fitted the two-peak transitions of the chimera by a three-state model (A → B → C, S10) (red lines in Figs. 5B and 5C). The initial activation from A to B accounts for the low-temperature transition, while the second step from B to C pertains to the high-temperature peak. The fitting resulted in an activation energy E_1_ = 76 ±5 kcal/mol (n=8) and an end-state enthalpy change ΔH_1_ = 96±9 kcal/mol (n=8) for A → B, and E_2_ = 115 ±3 kcal/mol and ΔH_2_ = 263±30 kcal/mol (n=8) for B → C, respectively. The total enthalpy change in the chimera was ΔH_total_ ∼ 359 ± 38 kcal/mol (n=8). Both the activation energies and the end-state enthalpy changes were profoundly reduced in the chimera as compared to those in the wild-type channel (Fig. 4d). For a multi-component process, an explicit mapping between calorimetric and functional measurements is challenging. Nonetheless, the chimeric changes of the DSC profiles were on par with functional changes. For example, the occurring of the low-temperature DSC peak closely match the large decrease of the temperature threshold of heat activation of the chimera ^26^. Furthermore, the low activation enthalpy E and end-state ΔH of the peak also mirrored the profound reduction of the temperature dependence in the chimera^24,26^. These observations suggest that the low-temperature DSC transition correlated with functional heat activation of the chimera. Consistently, the high-temperature peak occurred at temperatures above heat activation, and with an energy close to protein unfolding, it pertained to the major unfolding transition. Regardless of functional correspondences, the results provide explicit evidence about the multi-component composition of the DSC transition in wild-type channels and the impact of chimeric swap on the concertedness between the component transitions. The wild-type proximal N-terminus apparently plays an essential role in fortifying the concerted activation. As a control, the maltose binding protein fused to the chimera behaved the same as in the TRPV2-MBP (Fig. 4b insert), in good keeping with its independent thermal transitions.

We next probed the reversibility of the low-temperature transition of the chimera (Fig. 5). We first limited the first DSC scan to a final temperature 45 °C, before the onset of the high-temperature transition peak (Fig. 5d). While the low-temperature transition remained in the initial scan (Fig. 5a), it vanished in the subsequent, full-temperature scan (Fig. 5b). Instead, only the high-temperature transition was detected in the repeated scan (Fig. 5b), alongside the MBP peak (Fig. 5c). This experiment demonstrates that the chimera’s low-temperature transition, like the heat activity rundown, was irreversible even at relatively low temperatures (45 °C; Fig. S2e), suggesting unfolding was also involved. Biochemical experiments with native gel electrophoresis showed that the DSC sample proteins all belonged to the same tetrameric group (Fig. S3), supporting that both low- and high-temperature transition peaks originated from the same chimeric channel. The disruption of the concertedness of wild-type transitions appeared to be also specific to the swap of the MPD region. For comparison, we measured the DSC transition of a homolog TRPV1 channel lacking 15 residues in the turret region and the distal N terminus ^53^ (Fig. S4). Despite these significant changes, the mutant channel retained a single concerted, wild-type-like DSC transition. Consistently, the swap of the transmembrane domain (S1-S6) similarly retained a single concerted DSC transition in the chimera TRPV1/V2(S1-S6) (Fig. S5), while the transition temperature remained to correlate with the functional activation temperature ^26^, in agreement with the coupling of the opening and unfolding transitions.

The disruption of the coupling interactions between component transitions would leave the individual components to be more freely activated, in which case they may exhibit differential environmental sensitivity if different parts of channel proteins were involved. Thus, we repeated the DSC with the chimera in micelles. Fig. S6 shows the resultant thermograph of the chimera. Instead of two clearly separated transition peaks, the DSC transition in this case exhibited a single convoluted transition peak, with a noticeable hump on the rising phase (Fig. S6a). This thermograph thus also necessitates a multi-state mechanism. Furthermore, the hump occurred at similar temperatures to the low-temperature transition peak in liposomes, whereas the more prominent peak was shifted from 52 °C in liposomes to ∼45 °C in detergents (Figs. 5b vs. S6a). Thus, the change of the environment more prominently affected the unfolding transition. Quantitatively, the thermographs were adequately fit by a three-state model (A → B → C), resulting in E_1_ = 73 ±2 kcal/mol and ΔH_1_ = 98±11 kcal/mol for the initial activation, and E_2_ = 66 ±2 kcal/mol and ΔH_2_ = 262±11 kcal/mol (n=7) for the subsequent activation. The detergent environment impacted mainly the activation energy of the second transition, causing both the energy barrier and the temperature of the unfolding transition to be reduced. The two component thermal transitions in the chimera indeed possessed distinct propensity to environmental perturbations. Together, these results strongly support that the DSC transitions of the wild-type channels were a multi-component process involving distinct parts of the channel and that the MPD has an essential role in coupling them together for concerted activation. The findings suggest that the cooperative domain interactions are essential to driving up the energy barrier of activation and the temperature dependence of the channels.

## Discussion

This study on TRPV2 and its chimera aimed to provide direct molecular evidence supporting the suicidal model we recently proposed for TRPV1’s heat activation. Our findings identified specific molecular regions within the channels that regulate both heat activation and unfolding energy landscapes. The DSC of TRPV2 and its chimera also corroborated shared features between activation and unfolding, including similar activation energy barriers and transition temperatures, suggesting that channel opening triggers unfolding. This link between functional activation and unfolding is further evidenced by the concerted nature of transitions in wild-type channels, their resistance to environmental perturbations, specific temperatures requirements (Fig. 3), and independence from DSC heating protocols (Fig. 3). Thermal TRPs display unique temperature gating irreversibilities, such as heat-induced sensitization ^51^, hysteresis ^18^, incomplete closure ^26^, and rundown ^49^ (Fig. S7) – phenomena seldom seen in voltage- and ligand-dependent gating. These nonstationary behaviors, which have remained mechanistically elusive, may be explained by early unfolding events, further highlighting the inherent coupling of opening and unfolding. Supporting this, simulations incorporating such irreversible events reproduced the wild-type-like single concerted DSC transition despite their multi-state complexity (Fig. S8). Taken together, the coupling of functional activation and unfolding is supported by a diverse range of functional, molecular and calorimetric evidence.

The structural positioning of the proximal N-terminus in these channels provides further insight into the mechanisms of heat activation and unfolding (Fig. 6). TRPV2 and its homologs share membrane topology with voltage-gated channels ^54^. Immediately C-terminal to the pore-lining S6 lies the extended TRP helix, which runs parallel to the inner membrane and is flanked below by the proximal N-terminus and above by the cytosolic loops of the S1-S4 bundle, including the S4-S5 linker (L45). The interdomain interactions between the pore and both the S1-S4 bundle and the N-terminus (via TRP helix) create a resistive force to prevent channel gating under resting conditions. We hypothesize that elevated temperatures destabilize these interdomain interactions, either directly or through allosteric effects. In wild-type channels (Fig. 6a), effective channel opening requires concurrent destabilization of the S1-S4 bundle and N-terminus interactions, resulting in a high activation energy barrier and strong temperature dependence. The extensive rearrangement/breakdown of these interactions also initiates a cascading effect that further disrupt inter- and intra-domain interactions, eventually causing protein unfolding. In the chimera (Fig. 6b), replacing the proximal N-terminus appears to weaken its interactions with the TRP helix, (partially) decoupling the pore from N-terminal regions. This reduces the resistive forces and lowers the activation threshold and temperature dependence. The partial uncoupling also implies an uncoordinated destabilization of interactions. At lower temperatures, the chimera begins to open with less structural rearrangements, breaking down only a subset of interactions and leading to the initial, smaller DSC peak. The remaining channel structure remains stable and requires further heating to unfold, producing a subsequent high-temperature peak. Consequently, thermal transitions in the chimera tend to occur stepwise.

**Figure 6.**
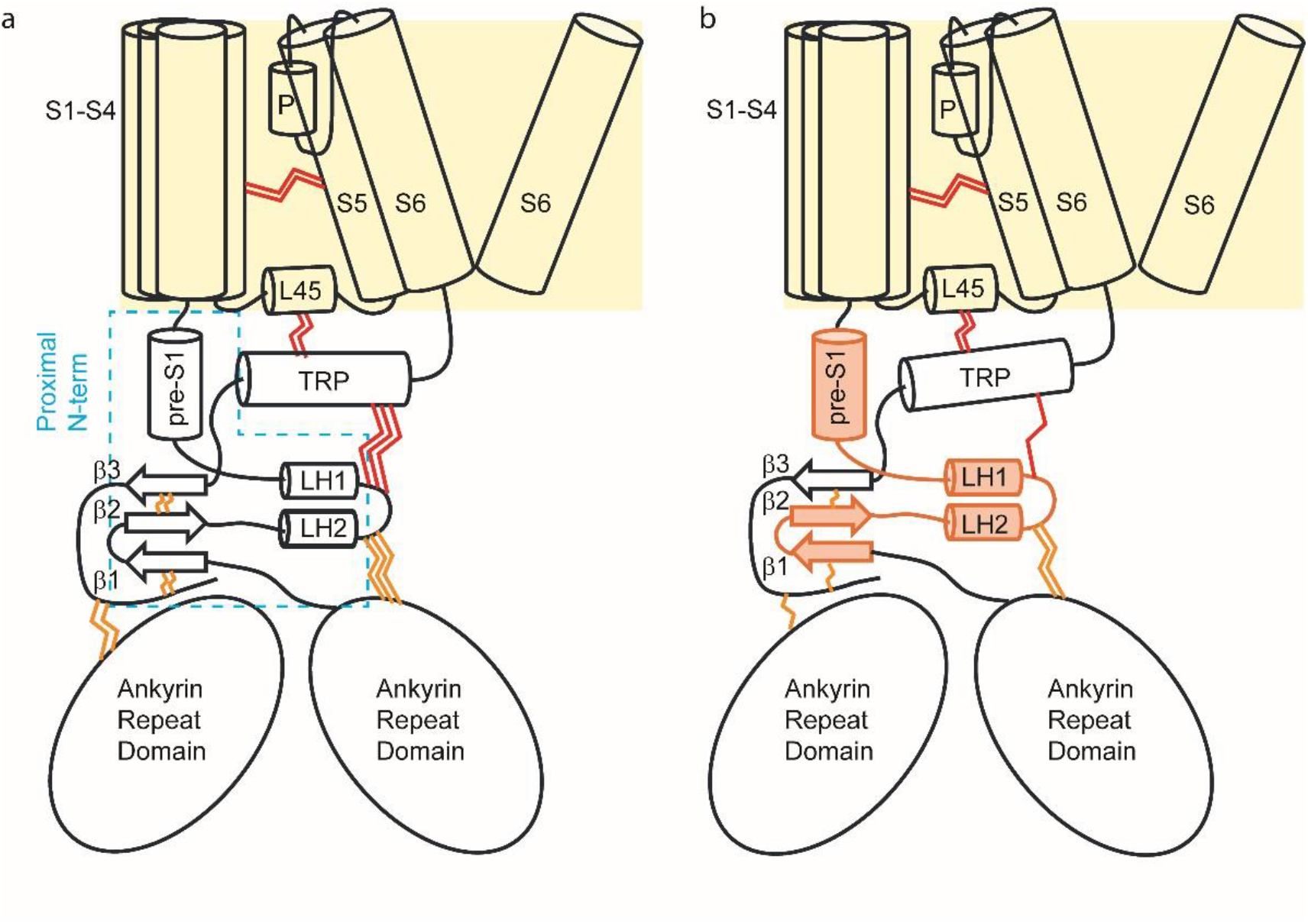
A conceptual model of heat activation and unfolding. **A)** A schematic of wild-type TRPV2 showing the pore (S5-S6) is interconnected with the S1-S4 bundle in the membrane and the N-terminus in the cytosol via the TRP helix at the C-terminus. Interdomain interactions (depicted by red wavy lines) exert a resistive force to prevent gate opening under resting conditions. Effective channel activation requires the simultaneous destabilization of both the S1-S4 bundle and the N-terminus interactions, resulting in a high activation energy barrier and a strong temperature dependence. Unfolding occurs due to an extensive rearrangement and breakdown of these interactions, initiating a cascade effect that ultimately leads to protein disintegration. **B)** In the chimera, replacing the proximal N-terminus (shown with orange filling) weakens interdomain interactions, partially decoupling the pore from N-terminal regions. This lowers the activation threshold and temperature dependence, making the chimera easier to activate while also experiencing less interaction breakdown, producing the initial smaller DSC peak. The remaining stable portion of the protein requires additional heating to unfold, resulting in the second DSC peak.

This proposed mechanism underscores the critical role of protein dynamics, particularly concerted interdomain interactions, in governing thermal sensitivity. The high temperature dependence of channel opening arises from the necessity to simultaneously overcome resistive forces from multiple domains. In retrospect, this model aligns well with various molecular findings on temperature gating. For instance, in the chimera with a modified proximal N-terminus, both the temperature dependence and activation threshold are reduced ^26^. This pattern diverges from the typical allosteric coupling in voltage- and ligand-gated channels, where weakening allosteric linkage generally reduces sensitivity while increasing activation threshold. Such mutagenic effects support the idea that interdomain interactions are directly involved in heat sensing. The molecular domains proposed to mediate these interdomain interactions – such as the N-terminal regions, the S1-S4 bundle, C-terminal regions, and pore domain –have all been implicated in temperature gating ^43-46^. Structural alterations in these regions can influence temperature sensitivity by modifying cross-domain interactions and their coordination. The unfolding model therefore offers a potential framework to unify these findings, which have been challenging to reconcile with other models. It is worth noting that most TRP channels are not temperature-activated, despite being prone to thermal unfolding (e.g. TRPV5 in Fig. S9 and ^52^). Therefore, thermal unfolding alone does not guarantee pore opening; activation occurs only if the enthalpic energy for pore opening is offset by entropic energy. Thermal TRPs, therefore, possess distinct thermodynamic properties that enable the coupling of unfolding to pore opening.

It may seem paradoxical that temperature receptors, designed to detect thermal fluctuations, become unstable at the very temperatures they are meant to sense. One possible explanation is that the strong temperature dependence of these channels demands substantial energy, making reversible conformational changes difficult to sustain. Intuitively, the more extensive the bond rearrangements in a protein, the lower the likelihood of fully reversible transitions. Indeed, as seen in thermal TRPs, irreversible changes are a common feature of temperature gating. The coupling of functional activation to unfolding not only explains this irreversibility but also posits that it is intrinsically tied to the heightened temperature sensitivity of these channels. This model predicts that similar irreversibility may also occur in other processes with strong temperature dependencies, providing a new framework for understanding the dynamic behavior of temperature responses.

## Materials and Methods

### 1. Expression and purification

Rat TRPV1 clone was provided by David Julius (UCSF). The BacMam expression vector was a gift of Erhu Cao (University of Utah). Protein expression and purification followed published protocols with modifications ^21^. Briefly, full-length channels were cloned into a BacMam expression vector ^55^ containing an N-terminal maltose binding protein (MBP) followed by a TEV protease site. Recombinant baculovirus encoding target genes were generated in SF9 cells using the Bac-to-Bac system (Invitrogen). The amplified P2 virus was used for infecting HEK293 GnTI^-^ cells, which were grown and maintained in Freestyle293 media (Gibco) containing 2.0 % FBS (Gibco) at 37 °C with constant shaking on an orbital shaker in a humidified incubator gassed with 8% CO_2_. Baculoviral transduction of cells took place at a cell density of ∼3.0 × 10^6^/ml. Infected cells were continuously incubated at 37 °C for ∼24 hrs and then shifted to 30 °C after addition of 10 mM sodium butyrate to boost protein expression. After ∼48 h of infection, cultures were harvested by centrifugation at 500 g for 15 min, and cell pellets were resuspended in a hypertonic buffer (36.5 mM sucrose, 50 mM Tris, 2.0 mM TCEP, pH8.0) supplemented with protease inhibitors (1.0 mM phenylmethanesulphonylfluoride [PMSF], 3.0 μg/ml aprotinin, 3.0 μg/ml leupeptin, 1.0 μg/ml pepstatin (all from Roche). The cell suspension was incubated on ice for 30 - 45 min in a Parr bomb at 500 psi followed by nitrogen cavitation. Cell lysates were clarified by brief low speed centrifugation to remove intact cells and nuclei. The membrane fraction was harvested by ultracentrifugation at 100,000 x g for 1 hr. The membrane pellet was resuspended in wash buffer (WB: 200 mM NaCl, 2.0 mM TCEP, 10% glycerol, 20 mM HEPES, pH 8.0) supplemented with protease inhibitors and solubilized by 1% DDM (Anatrace). The mixture was incubated at 4 °C for 1 hr with gentle shaking *prior to* centrifugation at 25,000 g for 30 min to remove insoluble fractions. The supernatants (membrane extracts) were collected and mixed with amylose resin (New England Biolabs). After 2 hrs of incubation at 4 °C, the resins were collected and transferred to a column which was washed with WB containing either 0.5mM DDM as well as 10 μg/ml soybean lipids (Avanti), followed by elution with 20 mM maltose in the same buffer. The purified sample was assessed for protein homogeneity by size-exclusion chromatography (SEC) in a Superose 6 column equilibrated with WB containing appropriate detergents and lipids and were used immediately for reconstitution experiments. Denaturing SDS-PAGE was performed on 8.0 - 10% polyacrylamide gels according to the Laemmli procedure using the Mini-PROTEAN Tetra System by Bio-Rad in a Tris/glycine/SDS running buffer (25 mM Tris, 192 mM glycine, 0.1% SDS, pH 8.3) and an applied voltage of 200 V. Native PAGE was performed on NativePAGE Bis-Tris precast gels (Invitrogen, Thermo Fisher Scientific) at 4°C following the manufacture’s protocol.

### 2. Protein reconstitution

Reconstitution of purified proteins into preformed liposomes was carried out using the detergent-mediated reconstitution approach ^21,56^. Lipids dissolved in chloroform were dried under a gentle nitrogen stream and then under a strong vacuum for 4-5 hrs before rehydration overnight in the reconstitution buffer at a concentration of 5.0 mg/ml (RB: 200 mM KCl, 20 mM MOPS, 2.0 mM TCEP pH 7). Liposomes were formed by 10 cycles of freeze-thawing and then 1-2 minutes of bath sonication. *Prior to* reconstitution, liposomes were destabilized by adding DDM to a final concentration of 4.0 mM, and the suspension was incubated at room temperature for 30 min with gentle shaking. Proteins were added to the detergent-lipid mixture at the desired lipid/protein ratio, typically 8:1 (weight/weight), and the mixture was incubated at room temperature for 1 hr under constant agitation. After equilibration, detergents were removed at room temperature using a fast detergent removal procedure through additions of four batches of prewashed polystyrene beads (Bio-Beads SM2, Bio-Rad) respectively at 30 mg, 30 mg, 50 mg and 100 mg per milligram of detergents. The first three treatments were incubated at room temperature at 1 hr intervals and the last one was at 4 °C overnight, all under gentle agitation. After complete detergent removal, the supernatant was aspired from BioBeads by a gel loading tip and then subject to ultracentrifugation at 100,000 g for 1 hr. The collected proteoliposomes were resuspended in the reconstitution buffer and flash frozen in liquid nitrogen for storage at -80 °C. For experiments requiring removal of the MBP tag from channel proteins, TEV was added to samples at a weight ratio of 20 : 1 (protein : Tev) before the last step of BioBeads addition.

### 3. Electrophysiology

Patch-clamp recording was carried out with an Axopatch 200B amplifier (Axon Instruments). Currents were low-pass filtered at 5–10 kHz through the built-in eight-pole Bessel filter and sampled at 10–20 kHz with a multifunctional data acquisition card (National Instruments). Data acquisition was controlled using custom software capable for synchronous I/O and simultaneous control of laser and patch-clamp amplifier. Patch pipettes were fabricated from borosilicate glass capillary (Sutter Instrument) with a resistance of ∼5 MΩ when filled with 150 mM NaCl solution. Pipette series resistance and capacitance were compensated using the built-in circuitry of the amplifier, and the liquid junction potential between the pipette and bath solutions was zeroed prior to seal formation. Currents were evoked from a holding potential at -60 mV.

Giant proteoliposomes suitable for patch-clamp recording were prepared using the dehydration-rehydration method as previously described ^21^. Briefly, frozen proteoliposomes (2-3 μl) were thawed and supplemented with 20 mM sucrose (final concentration). The mixture was dehydrated on a glass coverslip in a vacuum desiccator at room temperature (∼30 min), and then rehydrated with addition of ∼10 μl of reconstitution buffer (200 mM KCl, 5 mM MOPS, 2mM TCEP, pH 7) before overnight incubation at 4 °C in a sealed, humidified chamber. For patch-clamp experiments, the coverslip was transferred to a petri dish containing 150 mM NaCl, 2 mM MgCl_2_ and 10 mM HEPES (pH 7.4). Recordings were usually made on large unilamellar blisters at the peripheral of liposome clumps. The pipette and bath solutions were symmetrical. All chemicals were purchased from Sigma-Aldrich (St. Louis, MO).

Temperature jumps were produced by laser irradiation with infrared laser diode as previously described ^57^. In brief, the diode was mounted on a cooling block and operates at room temperature. Laser emission from the laser diode was launched into a multimode fiber with a 100-µm core diameter and 0.22 NA. The other end of the fiber was positioned close to patch pipettes as the perfusion pipette normally was. The laser diode was driven by a pulsed quasi-CW current power supply (Lumina Power). Constant temperature steps were generated by irradiating the tip of an open pipette and using the current of the electrode as the readout for feedback control. The laser was first powered on for a brief duration (<1 ms) to reach the target temperature and was subsequently modulated to maintain a constant pipette current. Between consecutive temperature pulses, laser power was adjusted manually and the adjustment generally took less than half a minute to a minute. Temperature was calibrated offline from the pipette current based on temperature dependence of electrolyte conductivity.

### 4. Calorimetry

DSC directly detects excess heat capacity changes by monitoring heat flow needed by a sample to keep up its temperature with that of the control buffer. DSC measurements were performed in a Microcal capillary ultrasensitive differential scanning microcalorimeter (Malvern Panalytical). Before each experiment, exhaustive cleaning of the cells was undertaken, and the sample and reference solutions were thoroughly degassed for at least 10-15 min in a temperature-controlled vacuum chamber (ThermalVac). About 250 ul degassed solution was loaded to each calorimetric cell, which was subsequently pressurized under an extra pressure of ∼2 atm to prevent solutions from degassing during heating. The measurements always started by filling both the sample and reference cells with the buffer used for the protein sample. The buffer-buffer scan was repeated until the instrument became thermally equilibrated, which typically took 4-5 repetitions. The sample was then loaded to the sample cell, normally during the cooling from the previous scan and before the start of thermal equilibration for the next scan. The sample-buffer scan was also repeated for 2-3 runs to determine the calorimetric reversibility of the thermally induced transition occurring during the first sample-buffer scan. Before the start of the first scan and between repeated scans, the solutions in the cells were allowed to equilibrate for 2-5 min at appropriate temperatures (4 or 15 °C for detergent-solubilized samples and 20 °C for reconstituted proteins).

For all measurements the scan rate was 1 °C/min, and protein concentrations were in the range of 0.4-0.8 mg/ml. The buffer used was either the reconstitution buffer for liposome experiments or the elution buffer of protein purification as described above. For solubilized proteins in detergents, the concentrations were determined from the absorbance at 280 nm using a Nanodrop spectrophotometer (ThermoFisher). For liposome samples, the concentrations were estimated based on the DSC transitions of the MBP tag fused to the channel proteins in reference to the pre-determined ΔH of MBP in detergents (∼220 kcal/mol, which is also similar to the published values of free MBP in aqueous solutions ^50^). For a rough estimation of protein concentration in liposomes (necessary for DSC sample loading), TRP fluorescence of the sample was measured in a fluorimeter (Hitachi F-7000), and the protein concentration was quantitated using BSA as a reference after subtraction of lipid background. The lipid recovery in our reconstitution assay was about 90%, as determined from fluorescence recovery by mixing 1% fluorescent lipids (ß-BODIPY FL C12-HPC, ThermoFisher) into reconstitution lipids. We also attempted more conventional quantitation methods such as BCA but with less accuracy.

### 5. Data analysis

The excess heat capacity thermographs were extracted from the original calorimetric profiles by first subtracting the repeated scan containing no transitions, followed by correction for possible baseline drifts as determined from pre- and post-transition baselines using established procedures as described in manufacturer’s user manual ^58^. The corrected profiles were normalized by protein concentrations to yield molar heat capacity. Equations for theoretical excess heat capacity were derived following ^59^ and are summarized in S10. Model fitting was carried out in a MATLAB program employing the built-in non-linear least-square solver implementing the Levenberg-Marquardt and trust-region-reflective algorithms ^48^.

## Acknowledgements

This work was supported by grant R01 GM132762 from the National Institutes of Health. The authors would like to thank Drs. Qiuxing Jiang, George Makhatadze and Zhengrong Yang for helpful discussions and Dr. Dinesh Indurthi for reading the manuscript and assisting with schematic creation.

---

**Figure S1.**
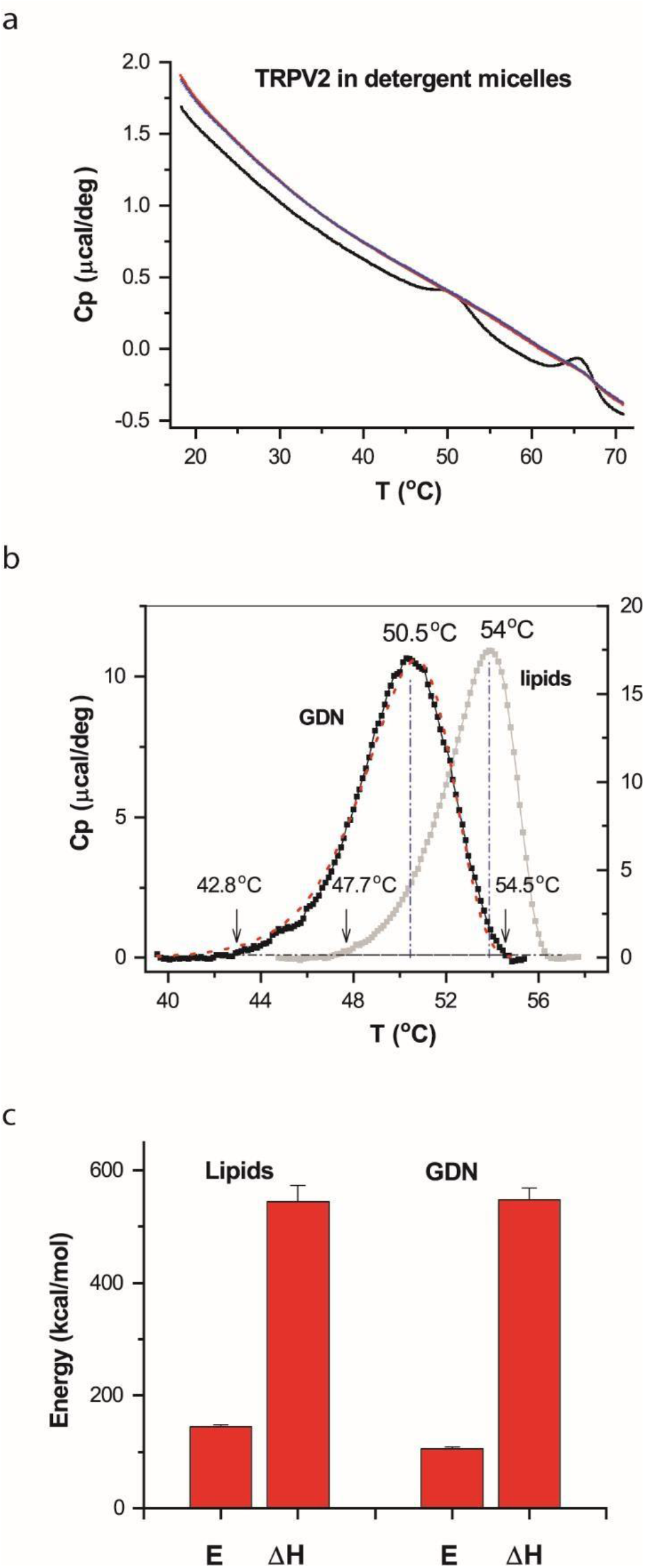
DSC transition of TRPV2 proteins in detergent micelles. Environmental perturbation of replacing lipids by detergent micelles produced a significant impact on transition temperatures, shifted from 54 °C to 50.5 °C, while individual component thermal transitions (e.g. channel opening, unfolding, etc) remained to occur as a single concerted peak. **A)** Representative DSC scans. Robust thermal transitions occurred during the first DSC scan but vanished in subsequent repeats. **B)** Thermographs of the excess heat capacity of TRPV2 proteins in detergent micelles (solid black; 0.02% glyco-diosgenin (GDN)) versus lipids (grey). The model fit was shown in red. Protein concentration was 0.44 mg/ml, determined from the absorbance at 280 nm. Shown in the background in grey color was the liposome measurement of TRPV2. The arrows and vertical blue lines labeled the temperatures corresponding to the peak or 1% of peak amplitudes. Compared with thermal transitions in liposomes, the transitions of TRPV2 in detergents was significantly shifted to lower temperatures, with temperature threshold becoming comparable to that of TRPV1. **C)** Statistic plot of energetic estimates of TRPV2 in detergents (n=5) versus in liposomes (n=9). DSC scanning rate was 1.0 °C/min.

**Figure S2.**
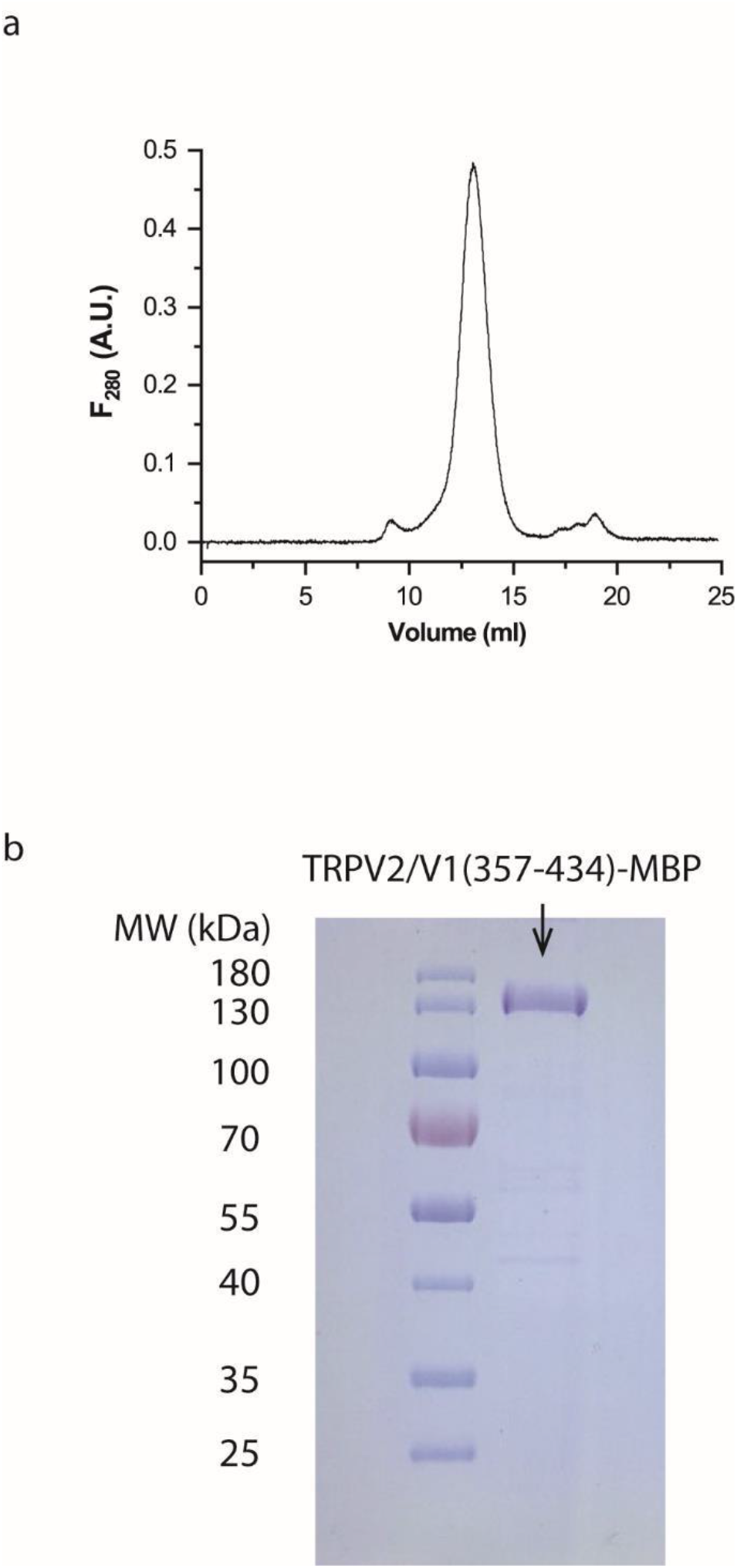
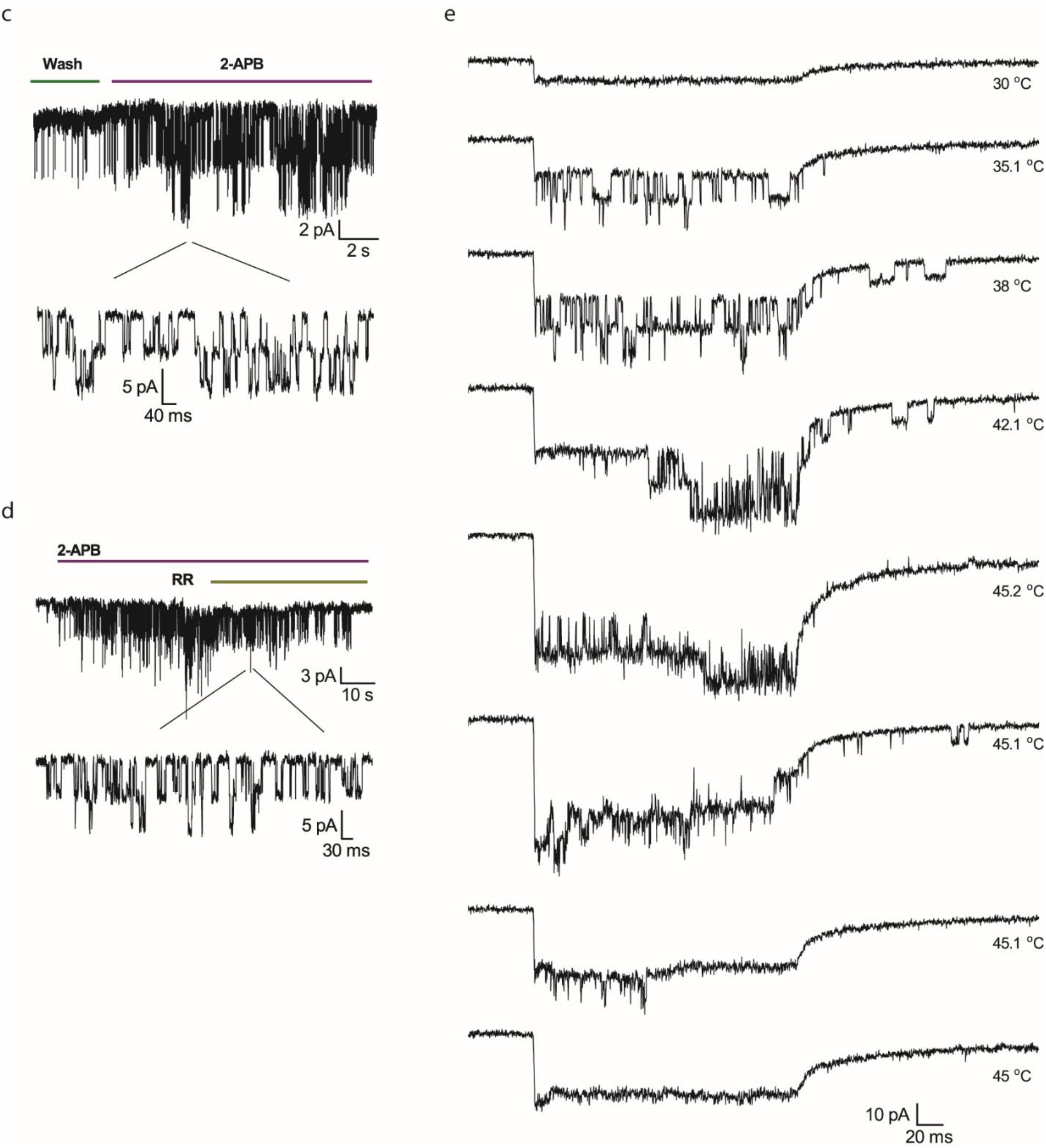
Characterization of chimeric TRPV2/V1(357-434)-MBP proteins. **A)** Size-exclusion chromatograph of purified chimeric proteins in a Superose 6 column. **B)** SDS-PAGE analysis of purified TRPV2/V1(357-434) by Coomassie blue staining. **C-E)** Patch-clamp recordings from reconstituted channels in liposomes, in response to 2-APB (1 mM; C), ruthenium red (D; 10 mM), and heat (E). Time between consecutive temperature pulses was on the order of seconds (time taken to adjust laser power). Continued heating caused the chimeric channel also to rundown as similarly happened to wild-type TRPV2. Holding potential -60 mV.

**Figure S3.**
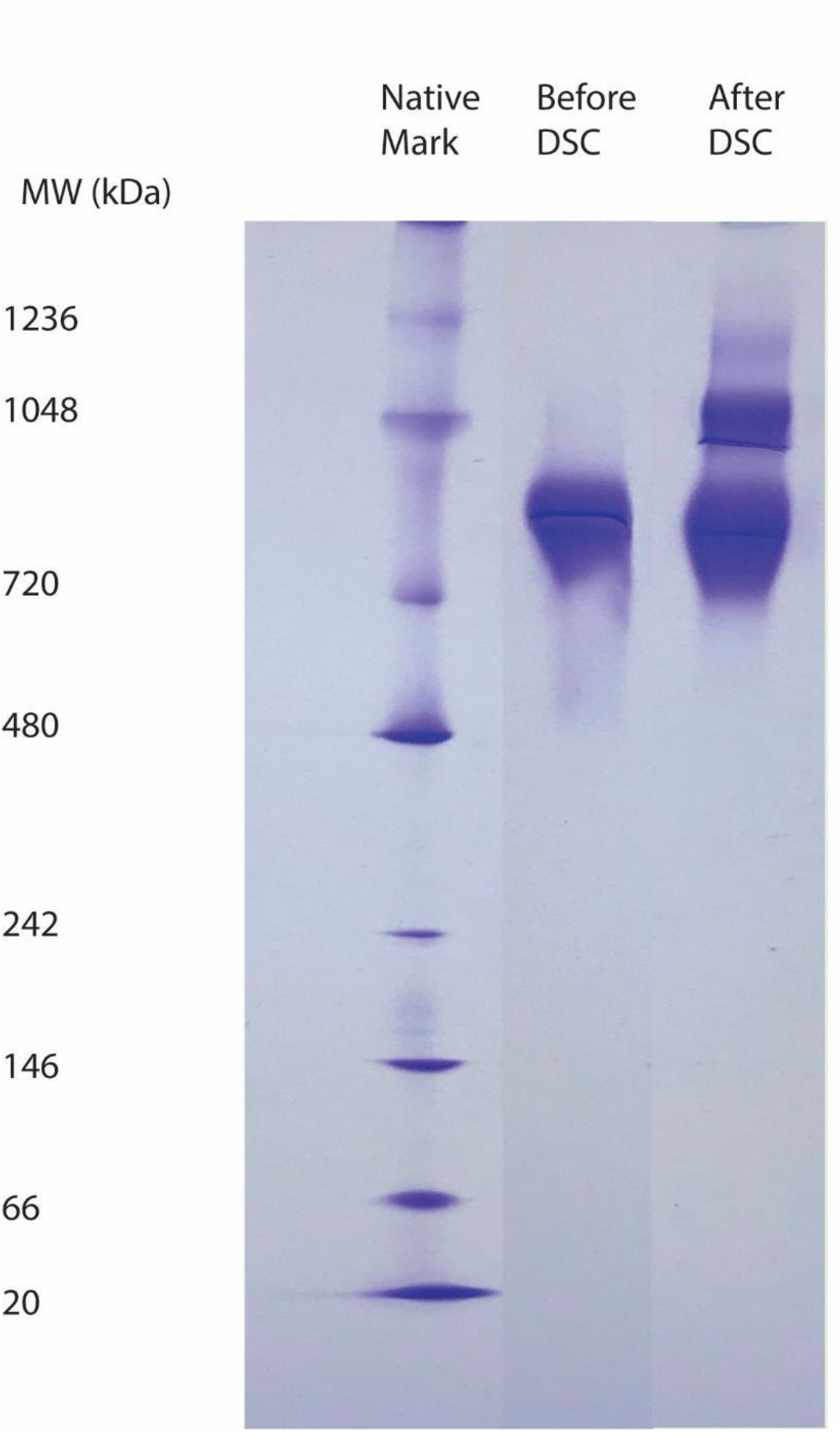
Native PAGE of chimeric channel TRPV2/V1(357-434)-MBP. Left lane: Native protein standards (Native-Marker). Middle: Purified, detergent-solubilized chimeric channels. Right: Same protein sample after DSC experiment at final temperature 55 °C.

**Figure S4.**
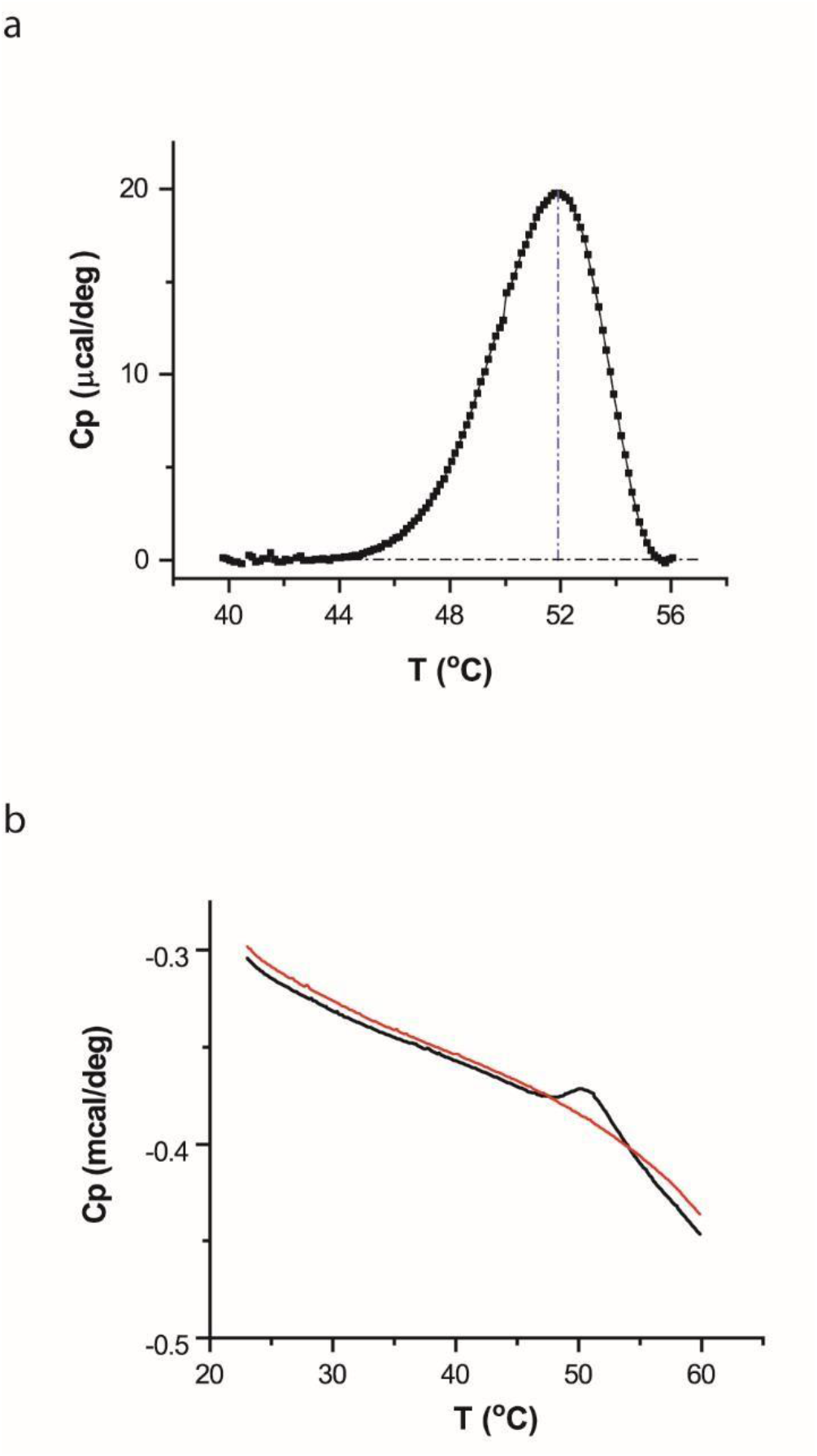
DSC profile of a TRPV1-mini channel with deletions in both the pore turret region and the distal N-terminus preceding the ankyrin repeats. **A)** Robust thermal transitions remained detectable in the mutant channel with an overall profile similar to that of the wild-type TRPV1. The red dotted line represents the model fit. **B)** Representative DSC scans (black: 1st san; red: repeat).

**Figure S5.**
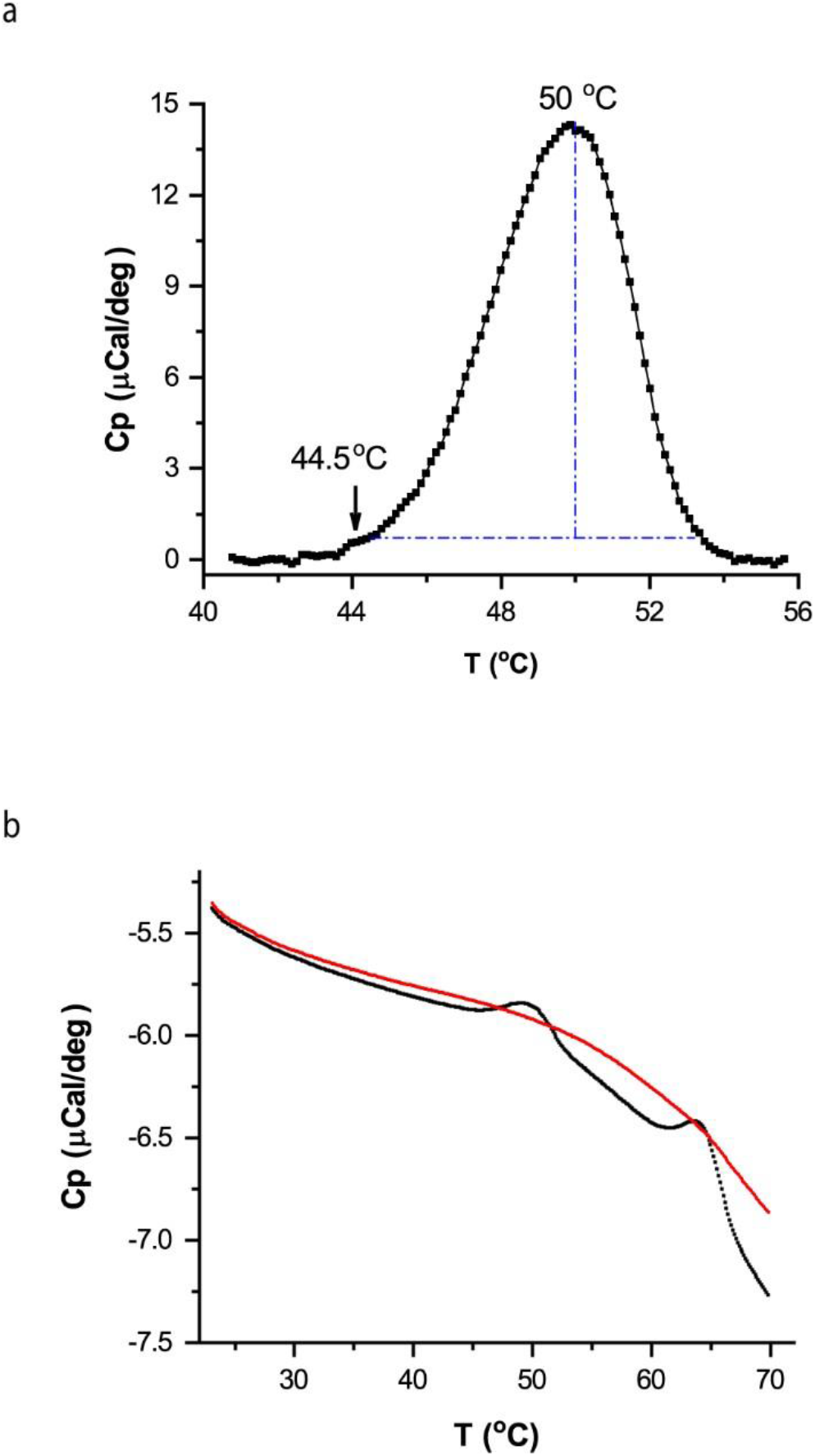
DSC of transmembrane domain chimera TRPV1/V2(s1-s6). **A**) Thermograph, showing a concerted transition peak retained after the exchange of transmembrane domain S1-S6. **B**) Representative DSC scan (black: 1^st^ scan, red: repeat).

**Figure S6.**
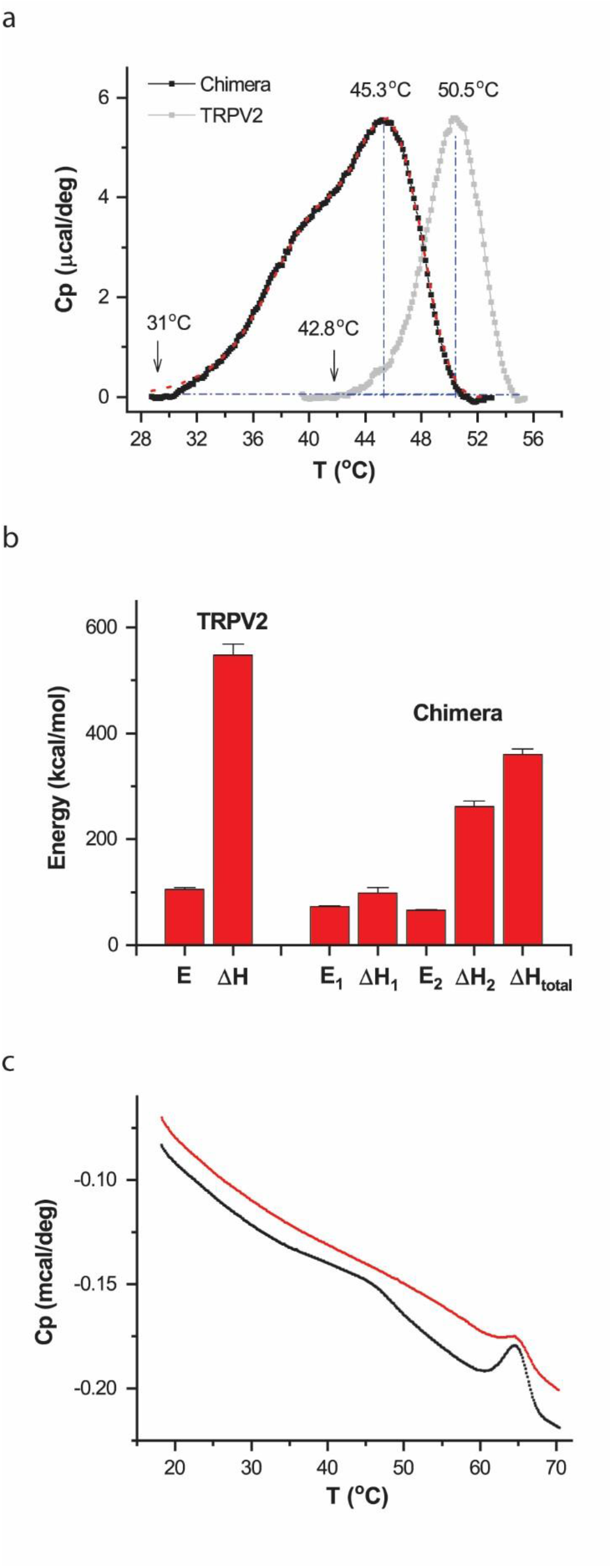
Thermal transitions of chimera TRPV2/V1(357-434) in micelles. **A)** Plot of excess heat capacity of TRPV2/V1(357-434) in micelles containing 0.02% glyco-diosgenin (GDN; black trace). The transition of the wild-type TRPV2 at the same conditions was shown in background in grey for comparison. The chimera shows a multitude of changes, including shift in temperature ranges, broadening of peak, and most noticeably, a hump occurring on its rising phase, which suggests that the transition also compromises two components as observed in vesicles. The arrows point to the temperature threshold at 1% of peak amplitude. The red line shows the fit by a three-state model (A→B→C). The first step of the model fits the hump on the rising phase, while the second step underlies the more prominent peak at higher temperatures. Protein concentration was 0.56 mg/ml, determined from absorbance at 280 nm. **B)** Summary plot of energetic estimates of thermal transitions (n=7), in comparison with those of wild-type TRPV2 in micelles. E_1_ and ΔH_1_ are respectively the activation energy and the end-state enthalpy change for the first transition (A→B), while E_2_ and ΔH_2_ correspond to those of the second step (B→C). ΔH_total_ represents the overall enthalpy change across all states. DSC scanning rate was 1.0 °C/min. **C)** Representative DSC scans. Black for initial scan, and red for repeat. DSC scanning rate

**Figure S7.**
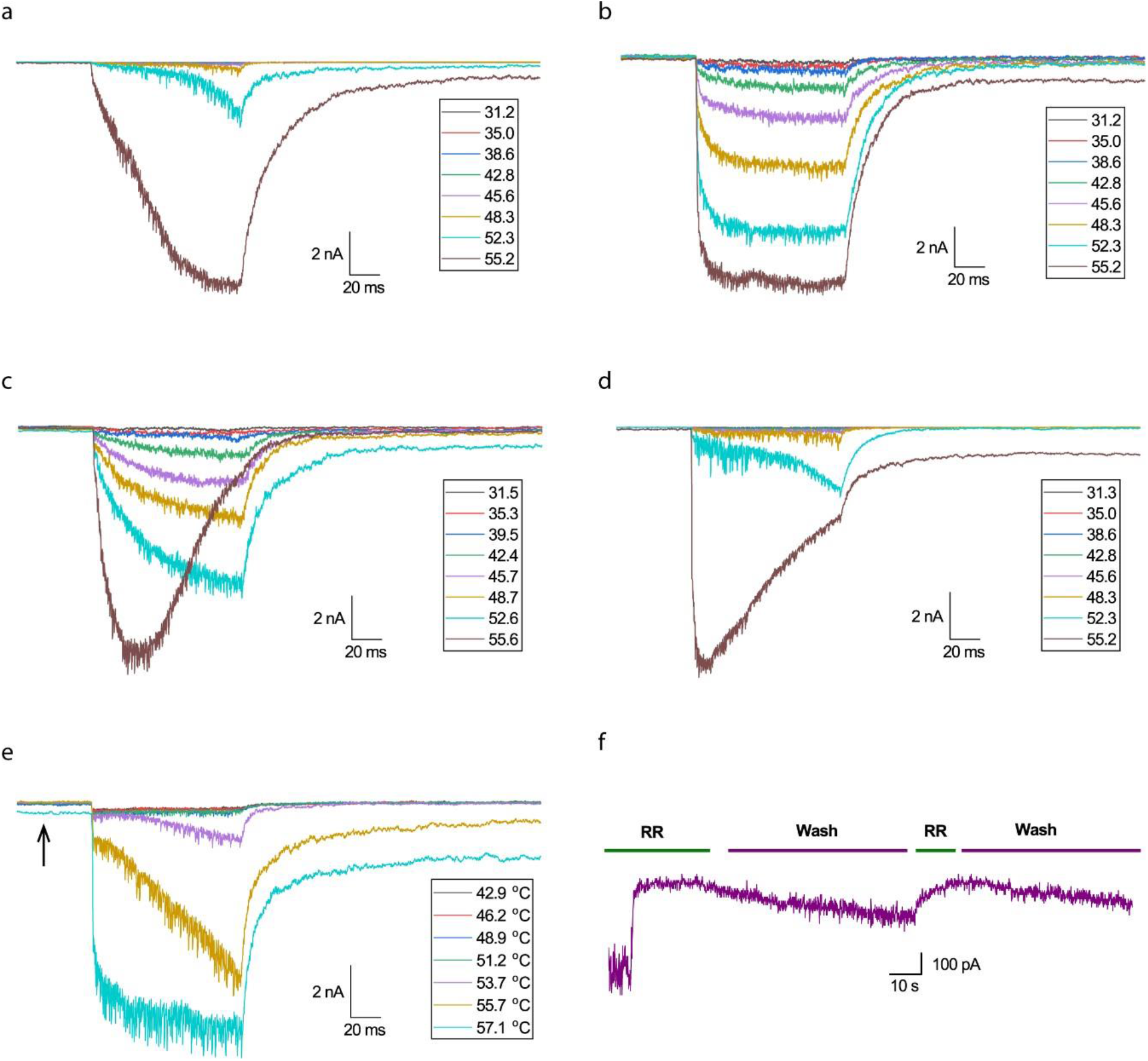
Heat responses of TRPV2, showing irreversible changes accompanying activation. Panels (*a*) and (*b*) illustrate sensitization. Initial responses (*a*) exhibited strong temperature dependence and high activation threshold, as opposed to responses during repeated run (*b*). The same family of temperature pulses were applied. Panels (*c*) and (*d*) show irreversible rundown, which could seemingly start at variable times. Responses (c) were obtained by sensitization. Panels (*e*) and (*f*) illustrate that the channel failed to shut off completely after heat activation ^26^. The arrow in (*e*) shows residual opening (leak) that was blocked by ruthenium red (RR), a pore blocker of TRPV2 (*f*). Recordings from HEK293 cells at -60 mV. Temperature pulses were 100 ms long, elevated from room temperature.

**Figure S8.**
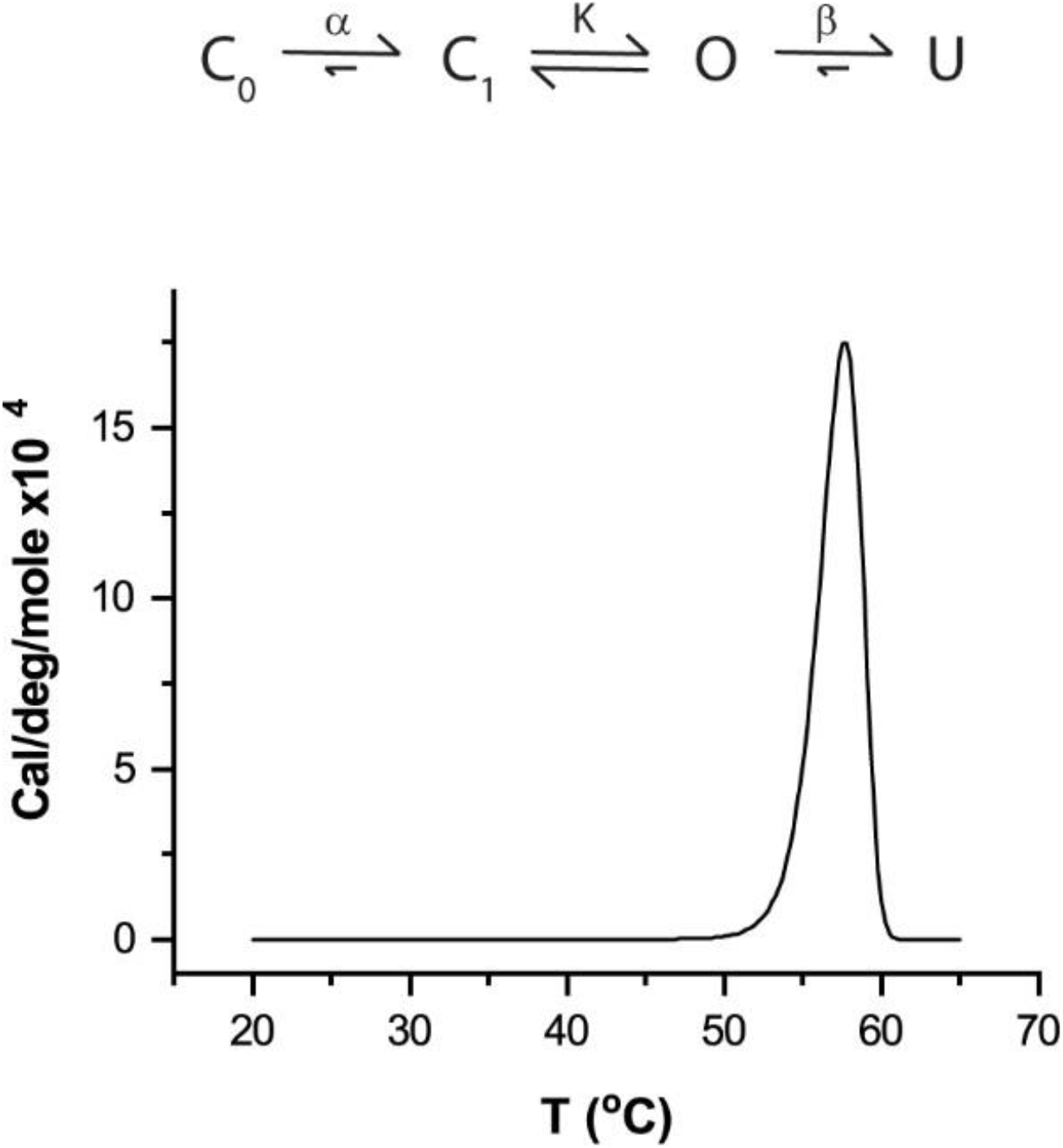
Illustration of single DSC transition from a multi-state model encapsulating sensitization, opening and unfolding. The model postulates that the channel resides initially in the resting state C_0_ and becomes irreversibly shifted to an intermediate closed state C_1_ upon heat stimulation. Transitions from C_0_ to C_1_ and from O to U (unfolded state) are irreversible and strongly temperature-dependent, whereas transitions between C_1_ and O are reversible and have moderate temperature dependence. Moreover, both irreversible transitions tend to occur at high temperatures with slow kinetics, whereas transitions between C_1_ and O at lower temperatures with fast kinetics. The sensitization reflects a mode shift of gating due to depletion of occupancy of C_0_. Plotted is an example of the excess heat capacity of the model, simulated using parameter values: ΔH_α_=200 kcal/mol, ΔH_β_=400 kcal/mol, ΔH_κ_=20 kcal/mol, E_α_=150 kcal/mol, E_β_=100 kcal/mol, T_α_=58 °C, T_β_=55 °C and T_κ_=40 °C. They determine the rate constants according to k=exp(-E/R(1/T-1/T^*\**^) where R is the gas constant, E is activation energy, and T^*\**^ an arbitrary reference temperature, and the equilibrium constant K=exp(-ΔH_κ_/R(1/T-1/T_κ_). Formalisms of the simulation were described in S10.

**Figure S9.**
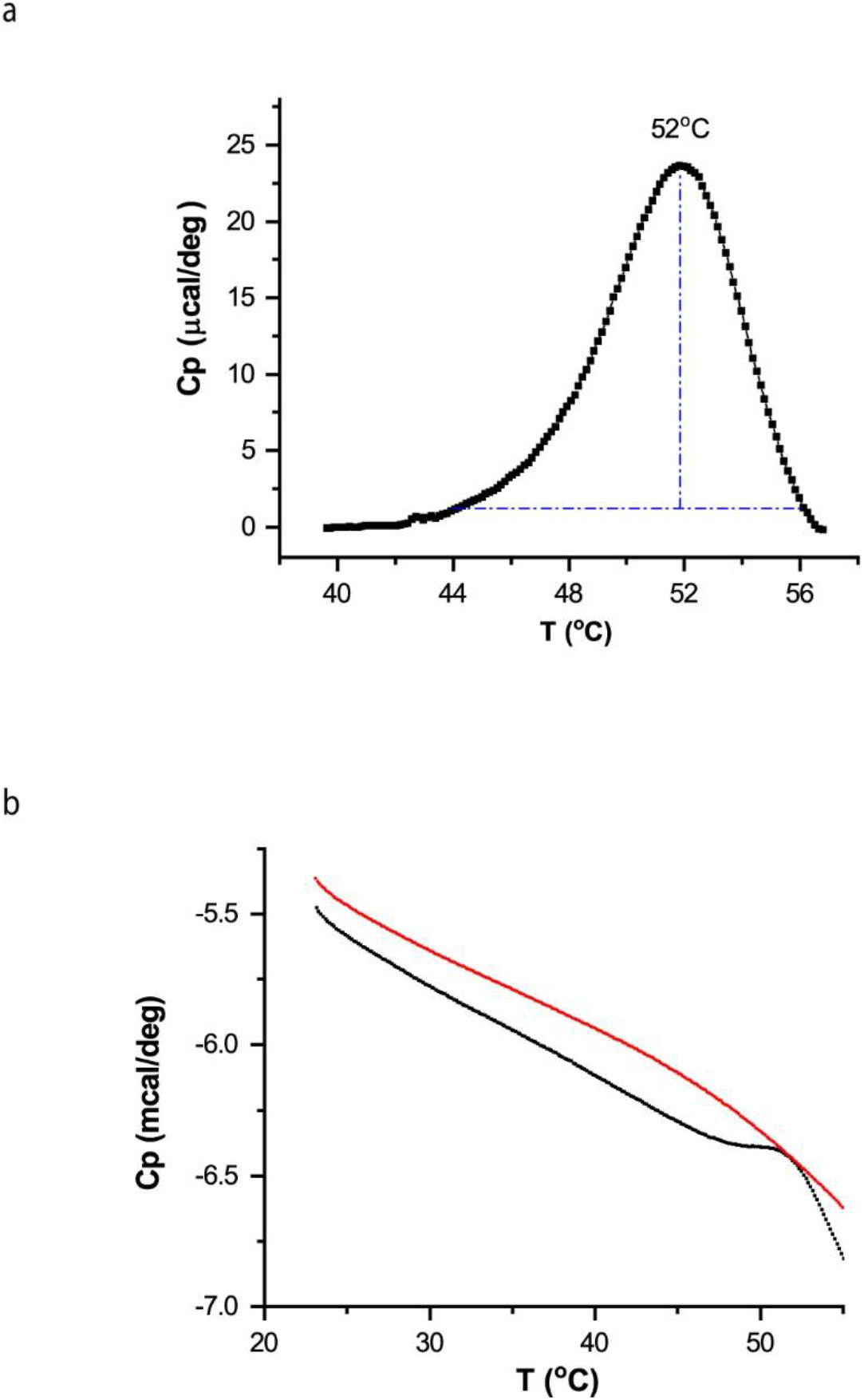
(**A**) DSC transition of TRPV5, showing robust unfolding transition at 40 °C - 55 °C, though the channel is not activated by heat. **B**) Representative DSC scans (black: initial, red: repeat).

## S10. Formalisms for the excess heat capacity of models used for data fitting

1. Two-state model:

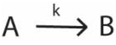

where *k* is the rate constant which is determined by the activation energy (E) according to:

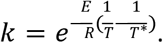

The excess heat capacity of the model follows:

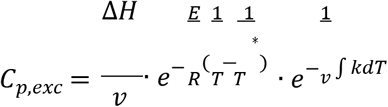

where v is the DSC scan rate.
2. Three-state model:

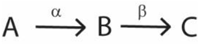

where α and β are rate constants and are related to their activation energies, E_1_ and E_2_ by

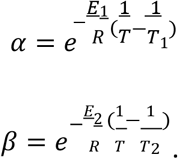

The excess heat capacity in this case can be derived as

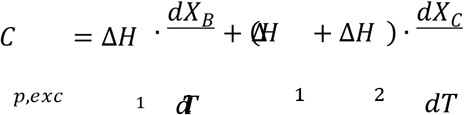

where ΔH_1_ and ΔH_2_ are enthalpy changes from A to B and B to C respectively, and X represents state occupancy and follows:

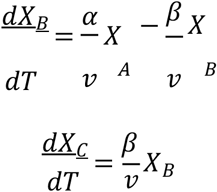

where v is the DSC scan rate. The excess heat capacity is solved by solving the above differential equations using Runge-Kutta formula with a trapezoidal rule (the ODE23TB solver in Matlab).
3. Four-state model:

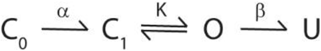

where rate constants α and β are related to their activation energies, E_α_ and E_β_ by

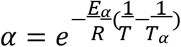

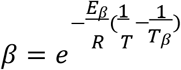

and equilibrium constant K depends on enthalpy change between C_1_ and O, ΔH_κ_:

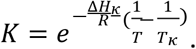

Let ΔH_α_ and ΔH_β_ be the enthalpy changes between C_0_ - C_1_ and O – U, and assume C_1_-O transitions at quasi-equilibrium. Then the excess heat capacity can be derived as

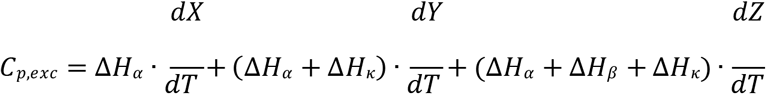

where X, Y and Z satisfy

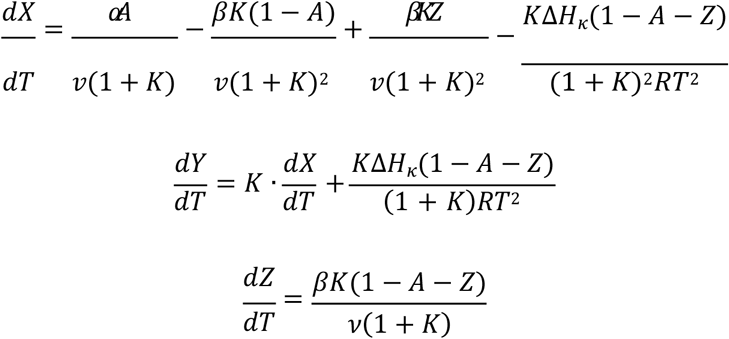

where 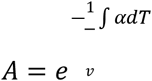 and v is the DSC scan rate. The excess heat capacity is solved by solving the above differential equations and integrals numerically in Matlab.

